# A novel Ca^2+^-binding protein influences photosynthetic electron transport in *Anabaena* sp. PCC 7120

**DOI:** 10.1101/571836

**Authors:** Julia Walter, Khaled A. Selim, Francisco Leganés, Francisca Fernández-Piñas, Ute C. Vothknecht, Karl Forchhammer, Eva-Mari Aro, Peter J. Gollan

**Affiliations:** Department of Biochemistry, Molecular Plant Biology, University of Turku, Tykistökatu 6A, 20520 Turku, Finland; Protein Evolution Department, Max Planck Institute for Developmental Biology, Max-Planck-Ring 5, 72076 Tübingen, Germany; Interfaculty Institute of Microbiology and Infection Medicine/Organismic Interactions Department, Eberhard-Karls-Universität Tübingen, Auf der Morgenstelle 28, 72076 Tübingen, Germany; Departamento de Biología, Facultad de Ciencias, Universidad Autónoma de Madrid, Calle Darwin 2, 28049 Madrid, Spain; Institute for Cellular and Molecular Botany, Plant Cell Biology, University of Bonn, Kirschallee 1, 53115 Bonn, Germany

**Keywords:** calcium, EF-hand, Anabaena, cyanobacteria, photosynthesis, phycobilisomes

## Abstract

Ca^2+^ is a potent signalling molecule that regulates many cellular processes. In cyanobacteria, Ca^2+^ has been linked to cell growth, stress response and photosynthesis, and to the development of specialist heterocyst cells in certain nitrogen-fixing species. Despite this, the pathways of calcium signal transduction in cyanobacteria are poorly understood, and only a few protein components are known. The current study describes a previously unreported calcium-binding protein which was called the Calcium Sensor EF-hand (CSE), which is conserved in filamentous, nitrogen-fixing cyanobacteria. CSE is shown to bind calcium, which induces a conformational change in the protein structure. Poor growth of a strain of *Anabaena* sp. PCC 7120 overexpressing CSE was attributed to diminished photosynthetic performance. Transcriptomics, biophysics and proteomics analyses revealed modifications in the light-harvesting phycobilisome and photosynthetic reaction center-protein complexes, and downregulated respiration.

## 1. Introduction

Calcium ions (Ca^2+^) play a pivotal role in a variety of cellular processes through its capacity to bind to proteins, changing their shape and charge [1]. A high intracellular Ca^2+^ concentration ([Ca^2+^]_i_) can be toxic because free Ca^2+^ can precipitate phosphate ions, which are essential for metabolic processes. Therefore [Ca^2+^]_i_ is closely monitored and tightly regulated by Ca^2+^ channels and pumps, and Ca^2+^-binding proteins [2,3]. In plants, free Ca^2+^ in the cytoplasm is maintained at around 100 nM, while higher concentrations are sequestered in various organelles. Rapid changes in the concentration of free Ca^2+^ encode signals that regulate numerous stress and developmental processes [reviewed in 4,5]. In the chloroplast, hormone signalling, photosynthesis and CO_2_ fixation are regulated by changes in Ca^2+^ concentration [reviewed in 6,7]. In cyanobacteria, which represent the photosynthetic ancestors of plant chloroplasts [8], Ca^2+^ was found to stimulate intracellular pH homeostasis in low external pH, thus preventing acidification of the cytoplasm and protecting physiological processes such as growth, photosynthesis and nitrogen (N) fixation from inhibition [9]. A Ca^2+^ signal observed in cyanobacteria during light-to-dark transitions [10], similar to the Ca^2+^ transient in plant chloroplasts that is induced by darkness [11], was partially inhibited by a Ca^2+^ channel blocker or calmodulin inhibitor. Hence, Ca^2+^ may be taken up in cyanobacteria from the extracellular medium by Ca^2+^-binding proteins or a Ca^2+^ pump, upon changes in the redox state of the plastoquinone (PQ) pool [10].

The major thylakoid protein complexes responsible for transformation of sunlight energy into chemical energy (ATP and NADPH) are well conserved between plants and cyanobacteria. Photosystem II (PSII), photosystem I (PSI) and the cytochrome b_6_f complex (cyt b_6_f) operate in series to transport electrons and to create a proton motive force that drives the ATP synthase complex. Ca^2+^ is essential for the water-splitting activity of the PSII oxygen-evolving complex [12–16], which liberates electrons from water to be used for CO_2_ fixation and concomitantly releases O_2_ to the atmosphere. PSII crystal structures of *Thermosynechococcus elongatus* and *T. vulgaris* showed that PSII monomers contain four Ca^2+^ ions, one of which is part of the oxygen-evolving complex and others are ligands in CP43 and PsbK [17,18]. PSI also binds calcium. Six Ca^2+^ ions were detected in the crystal structure of trimeric PSI in the *Synechocystis* sp. PCC 6803 PSI complex, two of which were localised to PsaL and PsaB subunits, suggesting a role in the oligomerisation of PSI or in providing a binding site for an unidentified regulatory protein [19].

While photosynthetic processes remain highly conserved between cyanobacteria and plants, one major difference concerns the capturing of photons. In plants, light energy is collected by the membrane-embedded light-harvesting complexes, which channel excitation energy towards the reaction centres of the photosystems. In cyanobacteria, however, light-harvesting antennae complexes, called phycobilisomes (PBS), are bound to the photosystems at the stromal face of the thylakoid membrane, with a special type of PBS being connected to PSI [20]. In *Synechocystis* sp. PCC 6803, even a megacomplex composed of PBS-PSII-PSI has been reported [21]. PBS are composed of different phycobilin pigment-binding proteins (PBP) and linker proteins, which are organised in allophycocyanin (APC) core cylinders connected to the photosystem reaction centres, and peripheral rod-shaped antennae linked to the APC core. The rods consist of core-connected phycocyanin (PC) discs, while some strains have additional phycoerythrocyanin (PEC) discs at the distal end of each rod [reviewed in 22]. Depending on the cyanobacterial species, the PBS composition and structure can differ in the number and type of PBPs, the type of bound chromophores (pigments) and the number of rods and core cylinders per PBS. In the multicellular model organism *Nostoc* sp. PCC 7120 (herein referred to as *Anabaena*), the APC core contains five cylinders from which eight rods radiate. APC, PC and PEC bind different numbers of the chromophore phycocyanobilin (PCB), resulting in different spectral features for each PBP. PEC absorbs light of shorter wavelengths (absorption maximum at 570 nm), whereas APC absorbs light of longer wavelengths (absorption maximum at 650 nm) [23]. Excitation energy is transferred from PEC via PC and APC to the terminal emitter ApcE, which is a pigmented core-membrane linker protein (L_CM_) connected to the PSII reaction centre in close proximity to CP43 [21,24,25]. Similarly, the terminal emitter ApcD connects PBS to PSI via hydrophobic interactions at the interface of two PSI monomers [20,21,26].Other linker proteins that do not bind pigments connect the discs, rods and cores within the PBS, to form complexes of around 6 000 kDa [24,27,28].

In this study, we present a previously undescribed EF-hand protein that is highly conserved in filamentous cyanobacteria. Based on demonstrations of Ca^2+^-binding, we called the protein “Ca^2+^ sensor EF-hand” (CSE). Over-expression of CSE in *Anabaena* was shown to affect photosynthetic electron transfer routes, leading to improper formation of light-harvesting PBS complexes and disrupting oligomerisation of the photosystems, which is essential for the connection of PBS to the reaction centres for functional electron transfer.

## 2. Materials and Methods

### 2.1 Growth conditions and treatments of Anabaena cultures

*Anabaena* cultures were grown in BG11_A_C medium, which is BG11_0_ supplemented with 10 mM NaHCO_3_ (BG11_0_C) [29] and 6 mM NH_4_Cl. CoCl_2_ ∙ 6 H_2_O was replaced with Co(NO_3_)_2_ ∙ 6 H_2_O, and medium contained 10 mM TES-KOH, pH 8.0, and. Cultures were grown under constant illumination of 50 μmol photons m^−2^ s^−1^ with gentle agitation (120 rpm) at 30°C in air enriched with 3% CO_2_. Liquid media for the the over-expressor strain (see below) included 1 μg ml^−1^ erythromycin. Fresh *Anabaena* cultures were started at an optical density (OD_750_) of 0.1 in BG11_A_C or BG11_0_C (for nitrogen-fixing conditions) as indicated. Total proteins and dry weight of cultures were measured as described in [30]. Chlorophyll *a* absorption (OD_665_) was measured in cultures suspended in 90% methanol, and pigment absorption spectra of cell suspensions were measured in whole-cell cultures with a Genesys 10S UV-Vis Spectrophotometer (Thermo Scientific) and a Shimadzu UV-1800 UV spectrophotometer (Berner), respectively.

### 2.2 Generation of Anabaena asr1131 over-expression and knockout strains

The *asr1131* gene sequence flanked by its native promoter and terminator sequences was amplified by PCR with the primers EF1 and EF4 (Table S1), cloned into the vector pST Blue and then into the *RSF1010*-based low-copy number plasmid *pRL1342* [31] using the *Xho*I and *Kpn*I restriction enzyme sites. This resulted in the overexpression plasmid *pBG2089* (see Figure S1), which was used for triparental conjugation of *Anabaena* wild-type (WT) as described in [32]. Transformants with overexpression of *asr1131* were selected on solid growth media containing 10 μg ml^−1^ erythromycin. Mutant strains of *Anabaena* lacking the *asr1131* gene were generated by replacing *asr1131* as well as non-coding DNA of 93 and 266 bp up- and downstream, respectively, with a neomycin/kanamycin-resistance cassette, and introducing the obtained plasmid via triparental conjugation. The *asr1131* knockout mutant strains were not thoroughly investigated in the current study (see Discussion).

### 2.3 Overexpression of recombinant protein

The *asr1131* coding sequence was amplified by polymerase chain reaction (PCR) using the oligonucleotides *cse*-NdeI-S and *cse*-EcoRI-AS (Table S1), and cloned into the pET-28a(+) vector (Novagen) for introducing a poly-His affinity tag. Recombinant His-tagged asr1131 protein was overexpressed in *Escherichia coli* (*E. coli*) BL21 cells grown at 37°C in Luria-Bertani (LB) medium supplemented with 50 μg ml^−1^ kanamycin by induction with 100 mM isopropyl β-D-1-thiogalactopyranoside (IPTG) and purified on a non-commercial Ni-NTA based affinity chromatography system.

### 2.4 ^*45*^*Ca*^*2*+^ overlay assay

Ca^2+^-binding capacity was tested in a radioactive ^45^Ca^2+^ overlay assay according to the method described in [33]. 20 – 80 μg proteins were spotted onto a methanol-activated PVDF membrane. The membrane was soaked three times in a Ca^2+^ washing buffer containing 60 mM KCl, 5 mM MgCl_2_ and 60 mM imidazole-HCl (pH 6.8) for 20 min at room temperature under gentle agitation. Afterwards, the membrane was incubated for 10 min at room temperature in the same buffer supplemented with 0.1 mM CaCl_2_ and 0.1 μM ^45^CaCl_2_ (13.90 mCi mg^−1^; Perkin Elmer), followed by 5 min rinsing with 50% ethanol. Subsequently, the PVDF membrane was completely dried between Whatman No. 1 filter paper and exposed overnight on a phospho-imaging screen. ^45^Ca^2+^ signals were detected with a FUJI FLA-3000 (FUJIFILM). Proteins were stained with Coomassie Brilliant Blue after the assay.

### 2.5 Isothermal titration calorimetry

Isothermal titration calorimetry (ITC) was performed in Tris-HCl buffer (pH 7.9), using a VP-ITC microcalorimeter (MicroCal) as described [34] after extensive dialysis of the purified recombinant protein in Chelex 100 (Sigma) to remove contaminating divalent cations. Calorimetric data were evaluated using MicroCal software (OriginLab) and fitted into one-site and two sequential binding sites models, for calculation of the binding thermodynamics [35]. ITC runs were repeated at least three times with two different batches of purified recombinant protein. From fitted curves, the association constants (*K*_*a*_) were generated and inverted to determine the dissociation constants (*K*_*d*_).

### 2.6 Size exclusion chromatography and multi angle light-scattering

Analytical size exclusion chromatography (SEC) using an ÄKTA chromatography system (GE Healthcare Life Sciences) fitted with a Superose 6 Increase 10/300 GL geometric column of 24 ml bed volume (GE Healthcare) was coupled to multi angle light-scattering (MALS) setup comprising a miniDawn Treos system (Wyatt Technology) and an Optilab T-rEX refractometer (Wyatt Technology). SEC MALS experiments were performed at room temperature using a flowrate of 0.5 ml/min, after equilibration of the column with 20 mM Tris-HCl buffer (pH 7.9), as indicated. The elution volume was plotted against the UV signal and the molecular mass was derived from the light scattering data. Data analysis and molecular weight calculations were done using ASTRA software (Wyatt) [36].

### 2.7 Circular dichroism spectra

Circular dichroism (CD) spectra of purified recombinant protein in 20 mM Tris-HCl buffer (pH 7.9) were recorded at 20°C from 195 to 250 nm using a J-810 spectropolarimeter (JASCO). Thermally-induced protein denaturation was determined by CD spectroscopy at 212 nm, recorded between 20 - 95°C. CD spectra of thermally-denatured proteins were recorded at 95°C, and protein refolding was recorded in samples subsequently cooled to 20°C [37].

### 2.8 Microscopy techniques

For the determination of the heterocyst frequency, cultures grown for two days in 3% CO_2_ in BG11_A_C or BG11_0_C (for nitrogen-fixing conditions) and cells were stained for 5 min using 0.5% Alcian Blue stain in 50% ethanol and washed three times with the respective growth media. Proheterocysts and heterocysts were counted from ×400 magnification micrographs taken with a Wetzlar light microscope (Leitz). 1000-2000 cells were counted for each treatment, and the heterocyst frequency calculated as a percentage of total cells counted.

### 2.9 Nitrogenase activity measurements

Nitrogenase activity was determined using the acetylene reduction assay described by [38]. 5 ml liquid cultures grown in 3% CO_2_ in BG11_A_C or BG11_0_C (for nitrogen-fixing conditions) were flushed with argon for 20 min and incubated in 23 ml vials with 10% acetylene in the headspace for 20 h, under 50 μmol photons m^−2^ s^−1^ light at 30°C with gentle agitation (120 rpm). 20 μl samples of headspace were analysed for ethylene content using a gas chromatograph (GC, Perkin Elmer Clarus^®^ 580) with a Carboxen 1010 PLOT 30 m × 0.53 mm capillary column and a flame ionisation detector (FID) using argon as a carrier gas. 1% ethylene was measured for calibration. The enzyme activity was calculated from the peak area and normalised to protein content.

### 2.10 RNAseq transcriptomics

Total RNA was isolated as described in [30] from four biological replicates of WT and *asr1131* over-expressor cultures grown for two days in BG11_A_C medium in 3% CO_2_. RNA samples were submitted to the Beijing Genomics Institute (Shenzhen, China) for single-ended library preparation and sequencing of RNA libraries using Illumina-HiSeq2500/4000. RNAseq reads were aligned with the Strand NGS 2.7 software (Agilent, USA) using the reference genome and annotations of *Nostoc* sp. PCC 7120, downloaded from Ensembl (EBI). The DESeq R package was used for normalisation and quantification of the aligned reads. Significantly differentially expressed genes were identified using a 2-way ANOVA test with Benjamini-Hochberg p-value correction for the calculation of the false discovery rate (FDR).

### 2.11 Photosynthetic fluorescence analysis

Cultures were grown for two days in 3% CO_2_ in BG11_A_C and adjusted to a chlorophyll *a* concentration of 7.5 μg ml^−1^ prior to measurements of low temperature (77K) and room temperature fluorescence emission spectra, chlorophyll *a* fluorescence and P700 absorbance, F_0_ rise, state transitions, light curves, single flash-induced fluorescence decay, and P700 oxidoreduction according to [39,40]. For fluorescence emission spectra, 5 μM Eosin Y was used as an internal standard at 536 nm for excitation with 440 nm light generated with a monochromator. Chlorophyll fluorescence and P700 absorbance were measured independently using 400 ms saturating pulses, with samples for all P700 measurements adjusted to 15 μg ml^−1^ chlorophyll *a*. The acceptor side limitation of P700 (Y(NA)) was calculated as (P_m_-P_m_’)/P_m_.

Oxygen evolution in light and oxygen consumption in darkness were measured with a Clark-type oxygen electrode (DW1, Hansatech) with constant stirring at 30°C. Cultures were adjusted to a chlorophyll concentration of 7.5 μg ml^−1^. Oxygen concentrations were recorded in darkness for 5 min before subsequent illumination with 400 μmol photons m^−2^ s^−1^.

### 2.12 Photosynthetic protein complex analysis and proteomics

Thylakoid membrane protein fractions were isolated from 2 day-old cultures according to [40]. 75 μg thylakoids were solubilised with 1.5% *n*-dodecyl-β-D-maltoside (DM) and separated by large pore blue native polyacrylamide gel electrophoresis (lpBN-PAGE) using an acrylamide concentration gradient of 3.5-12.5%. After lpBN-PAGE separation, gel strips were cut, and proteins denatured in Laemmli buffer containing 5% β-mercaptoethanol and 6 M urea at room temperature for 30 min. After solubilisation, the strips were laid onto a 12% SDS polyacrylamide gel containing 6 M urea, and proteins were separated in the second dimension and subsequently stained with Sypro Ruby Protein Gel Stain. For identification of proteins by mass spectrometry, protein spots were excised from SDS-PAGES gels and proteins subjected to in-gel trypsin digestion, as previously described [42]. Peptides were identified by nanoscale liquid chromatography/electrospray ionization tandem mass spectrometry (nLC/ESI-MS/MS) using a Q-Exactive instrument (Thermo Scientific). The MS/MS spectra were analysed using *Nostoc* sp. PCC 7120 annotations (GCA_000009705.1 downloaded from Cyanobase; http://genome.microbedb.jp/cyanobase/) using Proteome Discoverer v.2.2 (Thermo Scientific) as previously described [42].

### 2.13 Bioinformatics methods

The tertiary structures of translated gene coding sequences were predicted using the I-TASSER server (http://zhanglab.ccmb.med.umich.edu/I-TASSER/). Asr1131 homologs obtained from Basic Local Alignment Search Tool (BLAST) searches were aligned by Muscle in MEGA6. Gene sequences, identifications and descriptions were collected from CyanoBase (Kazusa Genome Resources; genome.microbedb.jp/cyanobase), KEGG (www.genome.jp/kegg/), UniProt (uniprot.org) and the National Center for Biotechnology Information (NCBI; ncbi.nlm.nih.gov) databases.

## Results

### A small ^Ca2+^-binding protein is highly conserved in filamentous cyanobacteria

*In silico* searches for Ca^2+^-binding domains in cyanobacterial genome databases identified a gene putatively encoding a protein with two Ca^2+^-binding EF-hand domains, which was previously not described in published literature. 15 homologs of the putative EF-hand protein were encoded in 13 cyanobacterial species, with two paralogs each occurring in *Calothrix* sp. PCC 7507 and *Aliterella atlantica* CENA595. The gene was found exclusively in filamentous cyanobacteria, with one exception being the unicellular species *Aliterella atlantica* CENA595, which is closely related to filamentous cyanobacteria [43,44]. Two of the filamentous species encoding the putative EF-hand protein were non-heterocystous (*Crinalium epipsammum* PCC 9333 and *Oscillatoria* sp. PCC 6506), although *Oscillatoria* sp. PCC 6506 is capable of fixing atmospheric N [45].

Alignment of amino acid sequences translated from all orthologues of the putative EF-hand protein (Figure 1A) demonstrated strong conservation of the N-terminal EF-hand domain, which typically comprises 12 amino acids that form a Ca^2+^-binding loop flanked on each side by nine amino acids forming α-helices [46]. The C-terminal EF-hand encoded in orthologues from *Crinalium*, *Hapalosiphon*, *Oscillatoria*, *Tolypothrix* and *Aliterella* displayed variation in the first five positions of the loop (positions 54 – 58 in Figure 1A), of which positions 1, 3 and 5 are known to be directly involved in binding of Ca^2+^ [46,47]. Orthologue sequences retrieved from genomes of several strains of *Anabaena, Nostoc*, *Calothrix* and *Cylindrospermum* possessed the conserved positions required for binding Ca^2+^ in both EF-hands. However, the α-helix preceding the second Ca^2+^-binding loop lacked three amino acids (between position 49 – 50 in Figure 1B), and the canonical hydrophobic residue preceding the first Ca^2+^ ligand was substituted with a tyrosine (see position 53 in Figure 1B).

**Figure 1.**
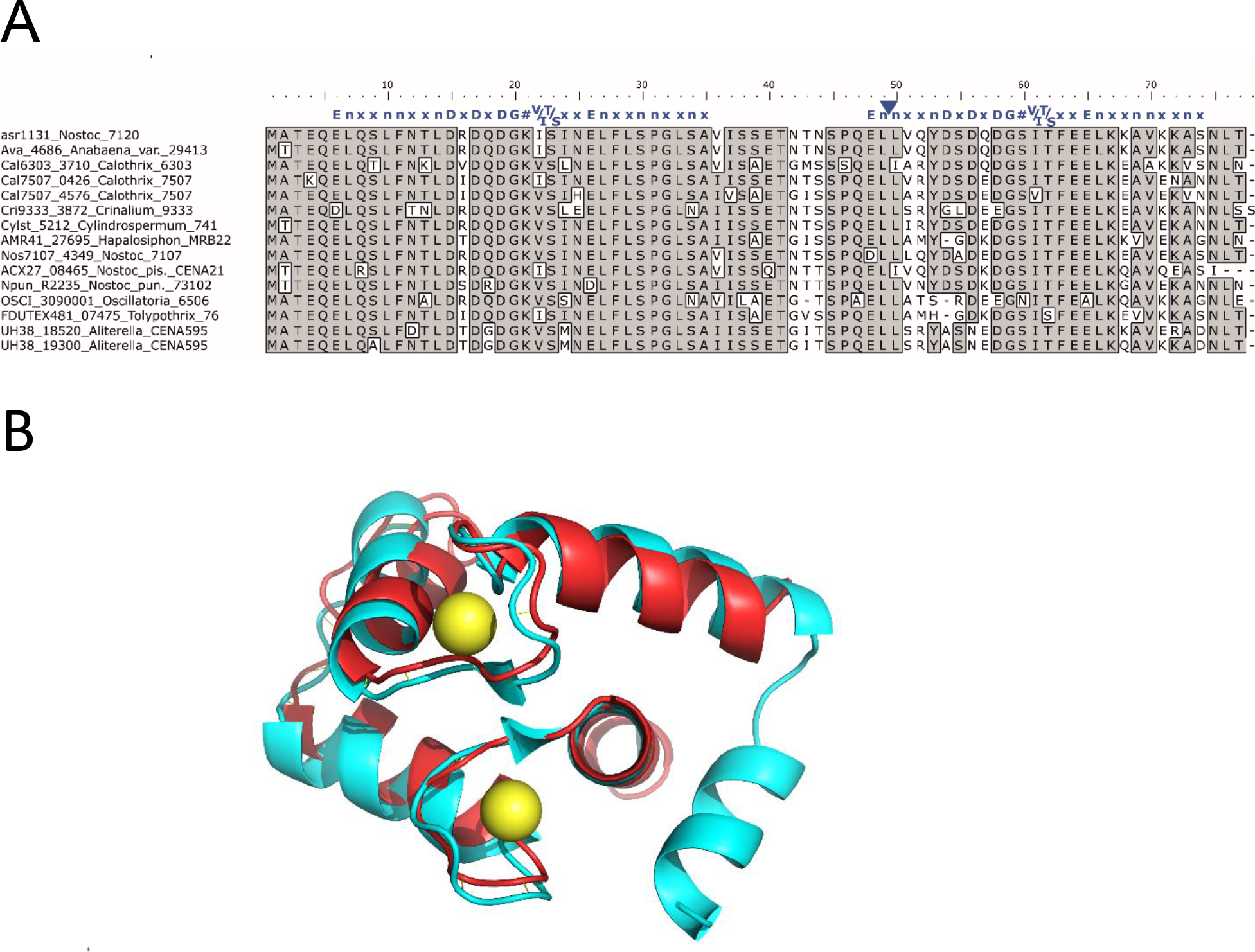
Multiple amino acid sequence alignment and predicted structure of putative EF-hand protein conserved in filamentous cyanobacteria **A.** Multiple sequence alignment of *asr1131* and other homologues from cyanobacteria. Identical residues are boxed and shaded. Canonical EF-hand sequence (blue letters) is aligned above the two EF-hand sequences of CSE, where specific amino acids in are represented by their single letter code, n is a hydrophobic residue, x is any residue, # is the fourth Ca^2+^ ligand (http://www.structbio.vanderbilt.edu/cabp_database/seq/indiv.aa/index.html). The blue triangle indicates a missing “xxn” motif from the second EF-hand **B.** Translated *asr1131* amino acid sequence (red) threaded through the crystal structure of the calcium-binding domain of mammalian troponin C (blue; PDB 2TN4) containing two bound Ca^2+^ (yellow);

In the filamentous cyanobacterium *Nostoc* sp. PCC 7120 (*Anabaena*) the putative EF-hand protein is encoded by *asr1131* and has a theoretical atomic mass of 8.5 kDa. Threading analysis to predict the tertiary structure of translated *asr1131* showed strongest resemblance with the helix-loop-helix structures of Ca^2+^-binding domains in calmodulin and troponin C (Figure 1B).

### 3.2 *In vitro biochemical properties of recombinant* asr1131 *protein*

The *asr1131* gene was cloned and over-expressed with an N-terminal His-tag. Purified recombinant protein was applied to size exclusion chromatography coupled with multiangle light scattering (SEC-MALS) for the determination of the oligomerisation state of recombinant asr1131 protein in the presence or absence of Ca^2+^. In Ca^2+^-Tris buffer, His-tagged asr1131 eluted as a monomer with an apparent mass of 9.65 kDa (Figure 2A). In Mg^2+^-Tris buffer lacking Ca^2+^, a minor shift in the elution profile of asr1131 was detected, corresponding to a molecular mass of 10.27 kDa, indicating a conformational change in the protein induced by Ca^2+^ ions (Figure 2A). The eluted peaks were collected and subjected to SDS-PAGE to confirm the presence of asr1131 protein in each fraction (Figure 2B). SEC-MALS analysis showed the protein to be monomeric in solution.

**Figure 2.**
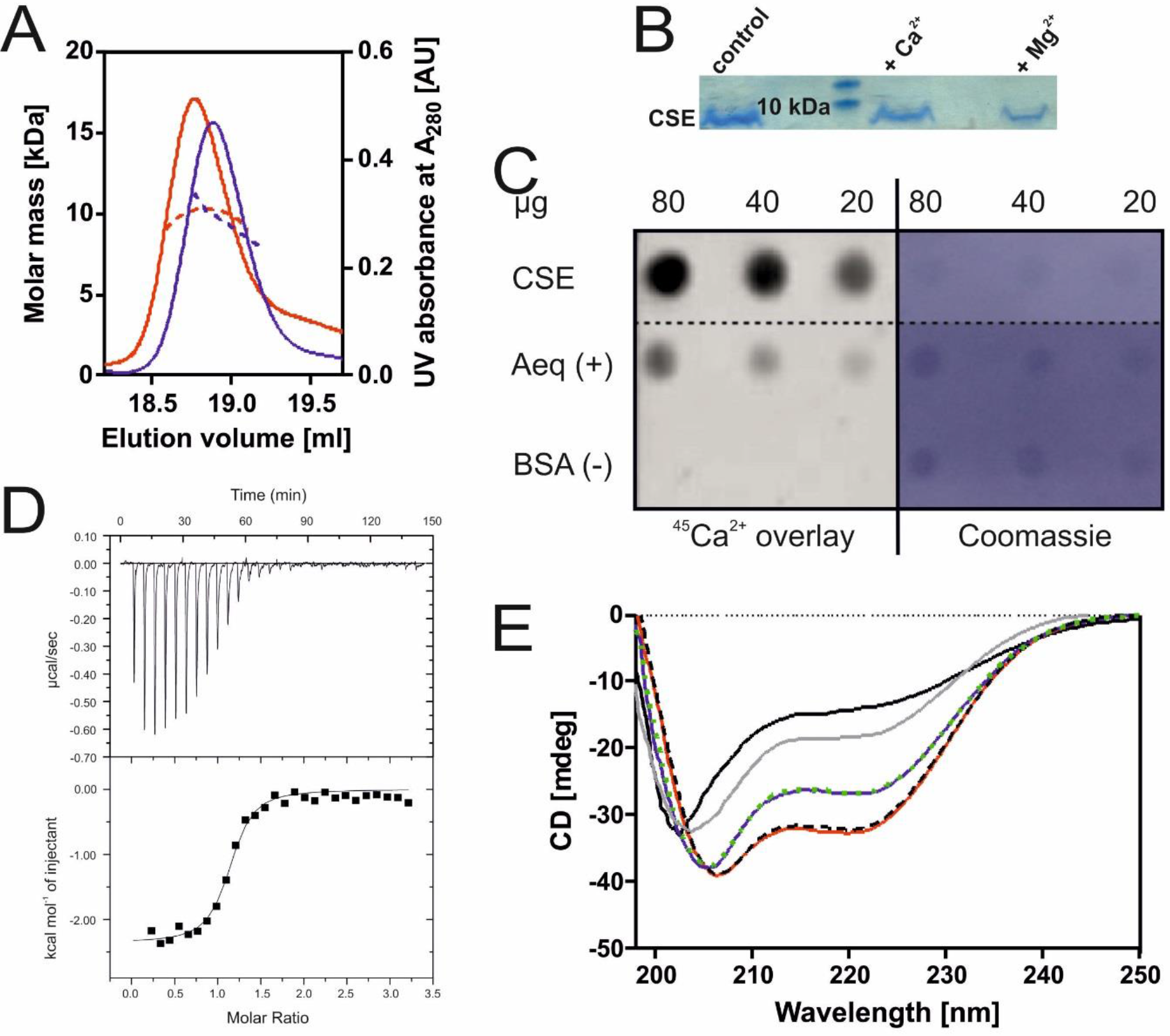
Structural analysis and Ca^2+^-binding properties of purified recombinant CSE protein **(A)** Size exclusion chromatography separation coupled to multiangle light scattering (SEC-MALS) used to detect the monomeric state of CSE in the presence of Ca^2+^ (blue line) and Mg^2+^ (red line), the dotted lines represent the distribution of molar mass across the respective peaks. **(B)** SDS-PAGE of collected protein peaks eluted from SEC-MALS; **(C)**^45^Ca^2+^ overlay assay of purified recombinant CSE protein. Aequorin (Aeq) and BSA were used as positive and negative controls, respectively. Image represents the results from three technical replicates. Proteins shown are from a single membrane that were treated together, and the image has been cropped to remove irrelevant proteins. **(D)** Isothermal titration calorimetry (ITC) metal-binding assay of recombinant His-tagged CSE (66 μM) in 1 mM CaCl_2_ in 20 mM Tris buffer. **(E)** Circular dichroism (CD) spectra at 20°C in different Tris buffers; without additions (black solid line); with NaCl (grey line); with NaCl and Mg^2+^ (blue line); with Mg^2+^ (green dashed line); with NaCl and Ca^2+^ (red line), with Ca^2+^ (black dashed line)

Three dilutions of purified recombinant His-tagged asr1131 were spotted onto a PVDF membrane that was subsequently overlaid with radioactive ^45^Ca^2+^. The autoradiogram showed efficient binding of Ca^2+^ to recombinant asr1131 *in vitro* (Figure 2C, top row). The weakest ^45^Ca^2+^ signal was obtained with the lowest protein amount (20 μg protein), and the signal intensity increased with each two-fold increase in protein amount. The ^45^Ca^2+^ signal obtained from recombinant asr1131 was even stronger than from the Ca^2+^-sensing photoprotein aequorin (AEQ), which was used as a positive control. Bovine serum albumin (BSA) used as a negative control showed no non-specific binding of ^45^Ca^2+^.

**Table 1.**
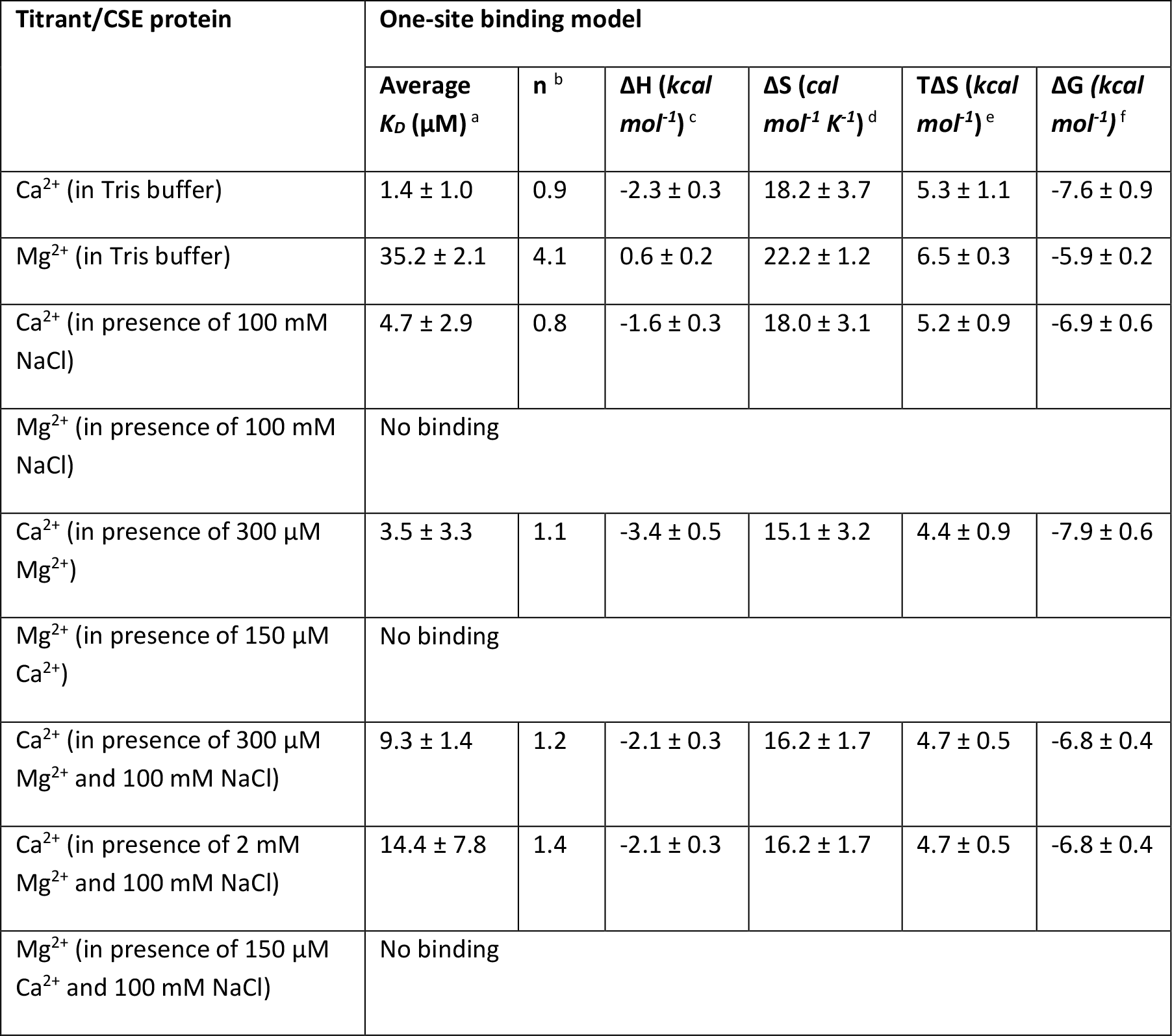
Thermodynamic parameters for Ca^2+^-binding to CSE protein The raw isothermal titration calorimetry (ITC) data were fitted using a one-site binding model for monomeric CSE. a The dissociation constant (*Kd*) values correspond to the mean of the independent experiments ± SD. All titrations were performed in 20 mM Tris-HCl based buffer (pH 7.9). ^b^ number of calculated binding sites ^c^ enthalpy ^d^ entropy ^e^ reversible heat ^f^ Gibbs energy

| Titrant/CSE protein | One-site binding model |  |  |  |  |  |
| --- | --- | --- | --- | --- | --- | --- |
| | Average $K_d$ ($\mu\text{M}$) <sup>a</sup> | n <sup>b</sup> | $\Delta H$ (kcal $\text{mol}^{-1}$) <sup>c</sup> | $\Delta S$ (cal $\text{mol}^{-1} \text{K}^{-1}$) <sup>d</sup> | $T\Delta S$ (kcal $\text{mol}^{-1}$) <sup>e</sup> | $\Delta G$ (kcal $\text{mol}^{-1}$) <sup>f</sup> |
| $\text{Ca}^{2+}$ (in Tris buffer) | $1.4 \pm 1.0$ | 0.9 | $-2.3 \pm 0.3$ | $18.2 \pm 3.7$ | $5.3 \pm 1.1$ | $-7.6 \pm 0.9$ |
| $\text{Mg}^{2+}$ (in Tris buffer) | $35.2 \pm 2.1$ | 4.1 | $0.6 \pm 0.2$ | $22.2 \pm 1.2$ | $6.5 \pm 0.3$ | $-5.9 \pm 0.2$ |
| $\text{Ca}^{2+}$ (in presence of 100 mM NaCl) | $4.7 \pm 2.9$ | 0.8 | $-1.6 \pm 0.3$ | $18.0 \pm 3.1$ | $5.2 \pm 0.9$ | $-6.9 \pm 0.6$ |
| $\text{Mg}^{2+}$ (in presence of 100 mM NaCl) | No binding | | | | | |
| $\text{Ca}^{2+}$ (in presence of 300 $\mu\text{M}$ $\text{Mg}^{2+}$) | $3.5 \pm 3.3$ | 1.1 | $-3.4 \pm 0.5$ | $15.1 \pm 3.2$ | $4.4 \pm 0.9$ | $-7.9 \pm 0.6$ |
| $\text{Mg}^{2+}$ (in presence of 150 $\mu\text{M}$ $\text{Ca}^{2+}$) | No binding | | | | | |
| $\text{Ca}^{2+}$ (in presence of 300 $\mu\text{M}$ $\text{Mg}^{2+}$ and 100 mM NaCl) | $9.3 \pm 1.4$ | 1.2 | $-2.1 \pm 0.3$ | $16.2 \pm 1.7$ | $4.7 \pm 0.5$ | $-6.8 \pm 0.4$ |
| $\text{Ca}^{2+}$ (in presence of 2 mM $\text{Mg}^{2+}$ and 100 mM NaCl) | $14.4 \pm 7.8$ | 1.4 | $-2.1 \pm 0.3$ | $16.2 \pm 1.7$ | $4.7 \pm 0.5$ | $-6.8 \pm 0.4$ |
| $\text{Mg}^{2+}$ (in presence of 150 $\mu\text{M}$ $\text{Ca}^{2+}$ and 100 mM NaCl) | No binding | | | | | |

Isothermal titration calorimetry (ITC) using His-tagged recombinant asr1131 protein was performed to define its Ca^2+^-binding stoichiometry and affinity. Raw ITC data were fitted into a one-binding site model to determine the stoichiometry of bound ligands (n), and the average dissociation constant (K_d_) for all available binding sites (Table 1, Figure S2). It was reported previously that NaCl can inhibit Ca^2+^-binding of the *Anabaena* Ca^2+^-binding protein CcbP [48], therefore the Ca^2+^- and Mg^2+^-binding properties of asr1131 were examined in the absence/presence of NaCl. In absence of NaCl, titration of asr1131 with ~5 μM CaCl_2_ per injection yielded exothermic calorimetric signals that saturated at a molar ratio of 1.11 (Figure 2D). Curve-fitting yielded a K_d_ of 1.4 ± 0.98 μM at a molar ratio of n = 0.94 (Table 1). These results corresponded to the binding of a single Ca^2+^ ion per asr1131 protein molecule. In contrast, the Mg^2+^-binding events showed an endothermic profile with a K_d_ value of 35.2 ± 2.1 μM, indicating very weak binding of Mg^2+^ (Table 1, Figure S2A), and supporting the specificity of asr1131 for Ca^2+^. Similar trends were exhibited in Tris buffer containing NaCl (Figure S2B); however, the binding enthalpy of Ca^2+^ was reduced (compare Figure 2D with S2B) and K_d_ values of 4.7 ± 2.9 μM were substantially higher than K_d_ in the absence of NaCl. Mg^2+^-binding was completely abolished by the addition of NaCl (Figure S2). Intracellular concentrations of free Mg^2+^ are much higher than Ca^2+^ *in vivo*, therefore Ca^2+^-binding by recombinant asr1131 under high Mg^2+^ were assessed. In 300 μM Mg^2+^, strong exothermic signals for each Ca^2+^ injection showed that Ca^2+^ bound to the asr1131 protein (Figure S2C), with a K_d_ value of 3.5 ± 3.3 μM and a molar ratio of n = 1.1 (Table 1) corresponding to binding of a single Ca^2+^ per protein molecule even in the presence of high Mg^2+^. Similar results were obtained in the presence of 100 mM NaCl, although with lower binding affinity of Ca^2+^ (K_d_ = 9.3 ± 1.4) (Table 1, Figure S2D). By contrast, the binding of 3 mM Mg^2+^ to asr1131 under 150 μM Ca^2+^ was completely suppressed, and Mg^2+^ was also unable to compete with Ca^2+^ for protein-binding in Tris buffer supplemented with 100 mM NaCl and 150 μM CaCl_2_ (Figure S2 and Table 1). Moreover, Ca^2+^ was able to induce strong exothermic binding signals and a K_d_ of 14.4 μM even in presence of 2 mM Mg^2+^ (Table 1, Figure S2E). Because the primary asr1131 sequence predicted two Ca^2+^-binding sites, the raw ITC data was also fitted to a model with two sequential binding sites, resulting in K_d_ values of 5.9 ± 7.5 and 56.3 ± 50.1 for the N-terminal and C-terminal binding site, respectively. Based on the ITC experiments, we concluded that under physiological conditions, recombinant asr1131 specifically binds one Ca^2+^ ion to the N-terminal binding site with high affinity, while the C-terminal site may have very low affinity to Ca^2+^ ions.

To identify the effects of Ca^2+^ on the protein secondary structure, the CD spectra of recombinant asr1131 were recorded in the presence or absence of Ca^2+^ and Mg^2+^. CD spectra exhibited typical α-helical profiles with two characteristic minima at 207 nm and 222 nm (Figure 2E). Ca^2+^ and Mg^2+^ caused a shift in the CD spectra towards increased α-helix peaks, indicating that both Ca^2+^and Mg^2+^ induced conformational changes in the protein, with Ca^2+^ inducing greater peak shifts than Mg^2+^. The melting curve of recombinant asr1131 protein from 20-95°C in the presence and absence of Ca^2+^ showed that a large proportion of the protein remained folded at 95°C (Figure S3A). Upon subsequent cooling to 20°C, the unfolded fraction of the protein regained the correct folding (Figure S3B), indicating that the asr1131 protein exhibits exquisite stability. Based on the Ca^2+^-binding activity and secondary structure determined here, we named the asr1131 protein “Ca^2+^ sensor EF hand” (CSE).

### 3.3 Overexpression of cse in Anabaena affects culture growth and pigment contents, but not heterocyst abundance or function

To explore the function of CSE *in vivo*, an *Anabaena* strain with upregulated expression of *cse* was engineered by transformation of wild-type *Anabaena* with the low copy number plasmid *pBG2089* (Figure S1) containing the *asr1131* gene under the control of its native promoter. This strain was found to exhibit two-fold higher expression of *cse* than WT *Anabaena* (Table 3). The *cse* over-expressor (*cse-*ox) strain was characterised in relation to WT *Anabaena*. Evaluation of culture growth rates over three days in CO_2_-enriched air (3% CO_2_), in BG11_A_C (N-replete) or BG11_0_C (N-fixing) media revealed about 30-50% slower growth of *cse-*ox in both media, compared to the WT (Figure 3A). Room temperature whole cell absorption spectra normalised to 750 nm showed a significant reduction in the PBS peak at 635 nm in *cse*-ox in both N-fixing and N-replete conditions (Figure 3B). The over-expressor also had a significantly lower chlorophyll peak at 685 nm, compared to the WT, when grown under N-fixing growth conditions. The carotenoid peak at 495 nm in the *cse*-ox was not significantly different from WT.

**Figure 3.**
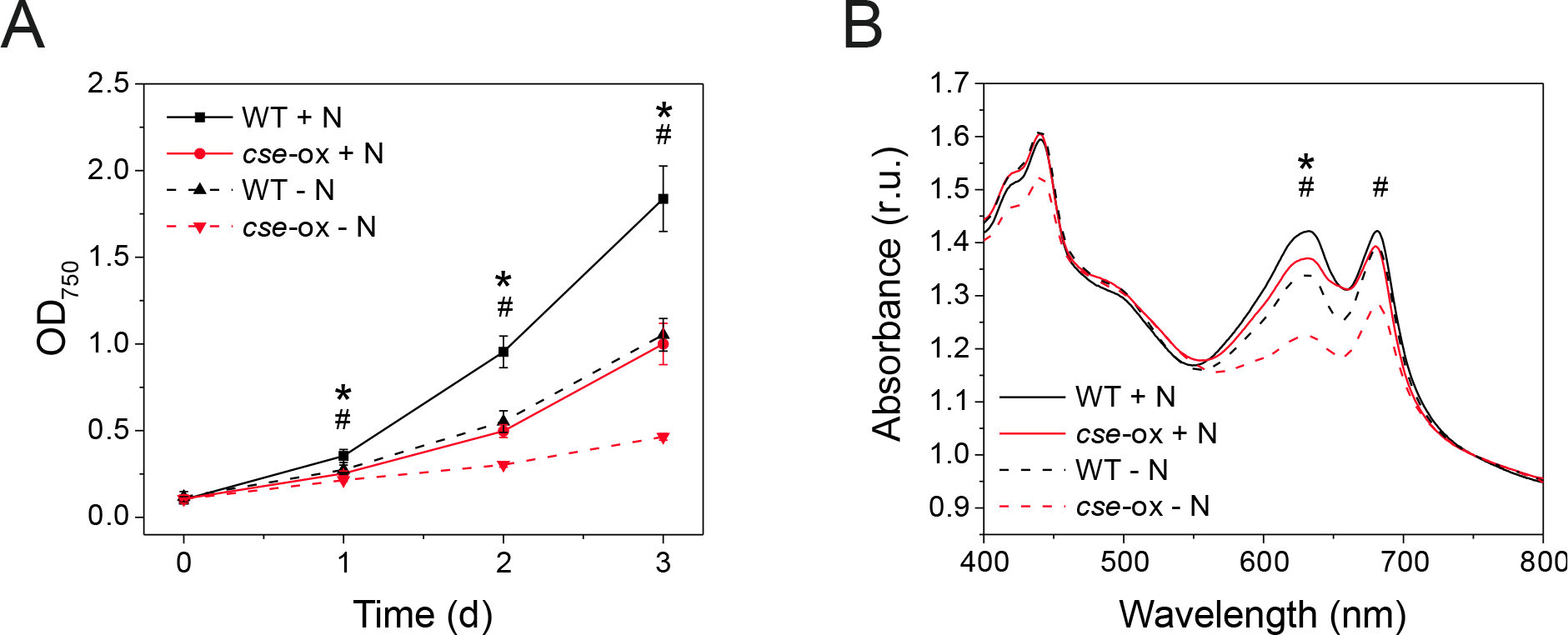
Growth phenotype induced by *cse* overexpression in *Anabaena* grown in nitrogen-replete and nitrogen-fixing conditions **(A)** Growth of *Anabaena* wild-type (WT; black) and *cse* overexpressor (*cse*-ox; red) strains monitored by absorbance at 750 nm. Cultures were grown in 3% CO_2_ in BG11_A_C medium (+N; solid lines) or BG110C (-N; dashed lines); **(B)** Absorption spectra of WT (black) and *cse*-ox (red) grown for two days in 3% CO_2_ in BG11_A_C medium (+N; solid lines) or BG110C (-N; dashed lines) normalised to absorbance at 750 nm. Peaks correspond to chlorophyll (440 nm and 680 nm), carotenoids (500 nm) and phycobilisomes (630 nm). Data points represent mean values from four biological replicates, error bars in **(A)** show standard deviations. Significant differences between WT and mutant samples are indicated with asterisks (+ N) and hashes (-N), respectively, (*t*-test *P*<0.05)

Comparisons of the protein and chlorophyll contents, the dry weight, heterocyst frequency and nitrogenase activity showed no significant differences between WT and *cse*-ox in N-replete conditions (Table 2). However, *cse*-ox cells grown for 2 days in N-fixing conditions had about 30% less chlorophyll and were around 12% heavier than WT cells. Heterocyst frequency and nitrogenase activity did not differ significantly between the two strains.

**Table 2.** Physiological parameters in N-replete and N-fixing conditions 2 day-old *Anabaena* WT and *cse* overexpressor (*cse*-ox) cultures were grown in 3% CO_2_ in BG110C medium either supplemented with NH_4_^+^ (+N) or lacking a source of combined N (-N). Errors indicate standard deviations that were calculated from three biological replicates, significant differences between WT and mutant samples are indicated with asterisks (*t*-test *P*<0.05)

| Strain and condition | Protein content (μg/ml) per OD<sub>750</sub> = 1.0 | chl content (μg/ml) per OD<sub>750</sub> = 1.0 | Dry weight (mg/ml) per OD<sub>750</sub> = 1.0 | Heterocyst frequency (% of total cells counted) | Nitrogenase activity (μmol h<sup>-1</sup> mg proteins<sup>-1</sup>) |
| --- | --- | --- | --- | --- | --- |
| WT + N | 273.1 ± 23.6 | 9.1 ± 0.3 | 0.46 ± 0.01 | 1.3 ± 0.5 | n.d. |
| *cse-ox* + N | 304.6 ± 11.3 | 9.1 ± 0.3 | 0.49 ± 0.02 * | 1.6 ± 0.3 | n.d. |
| WT - N | 218.5 ± 9.0 | 9.3 ± 0.2 | 0.44 ± 0.01 | 9.4 ± 1.0 | 2.49 ± 0.09 |
| *cse-ox* - N | 208.5 ± 10.1 | 6.3 ± 0.2 * | 0.50 ± 0.01 * | 9.3 ± 1.1 | 2.99 ± 0.37 |

### 3.4 Expression of genes encoding photosynthetic light-harvesting proteins is differentially regulated by cse overexpression

The transcriptomes of *Anabaena* WT and the *cse-ox* strain grown in BG11_0_C + NH_4_^+^ media in 3% CO_2_ were analysed after two days of growth in fresh medium. Significant differences were identified in expression of genes encoding subunits of the light-harvesting PBS complexes in *cse-ox*, compared to WT (Table 3). Members of the *cpc* gene cluster (*alr0528* – *alr0536*) encoding PC rods, and the entire *pec* cluster (*alr0523*–*alr0527*), encoding PEC rod caps [24] were strongly downregulated. Three genes encoding OCP-like N-terminal domain proteins (*all1123*, *all3221*, *alr4783*) were downregulated in the *cse*-ox, along with photosynthetic complex subunits *psaB* and *ndhD1* and several genes from the porphyrin pathway including *hemK*, *por*, *chlL* and *chlN*. Conversely *hemF2*, which is also involved in chlorophyll and porphyrin biosynthesis, was strongly upregulated in *cse-*ox compared to WT. Several genes involved in Ca^2+^-related processes were also differentially expressed in the over-expressor (see Table 3).

**Table 3.**
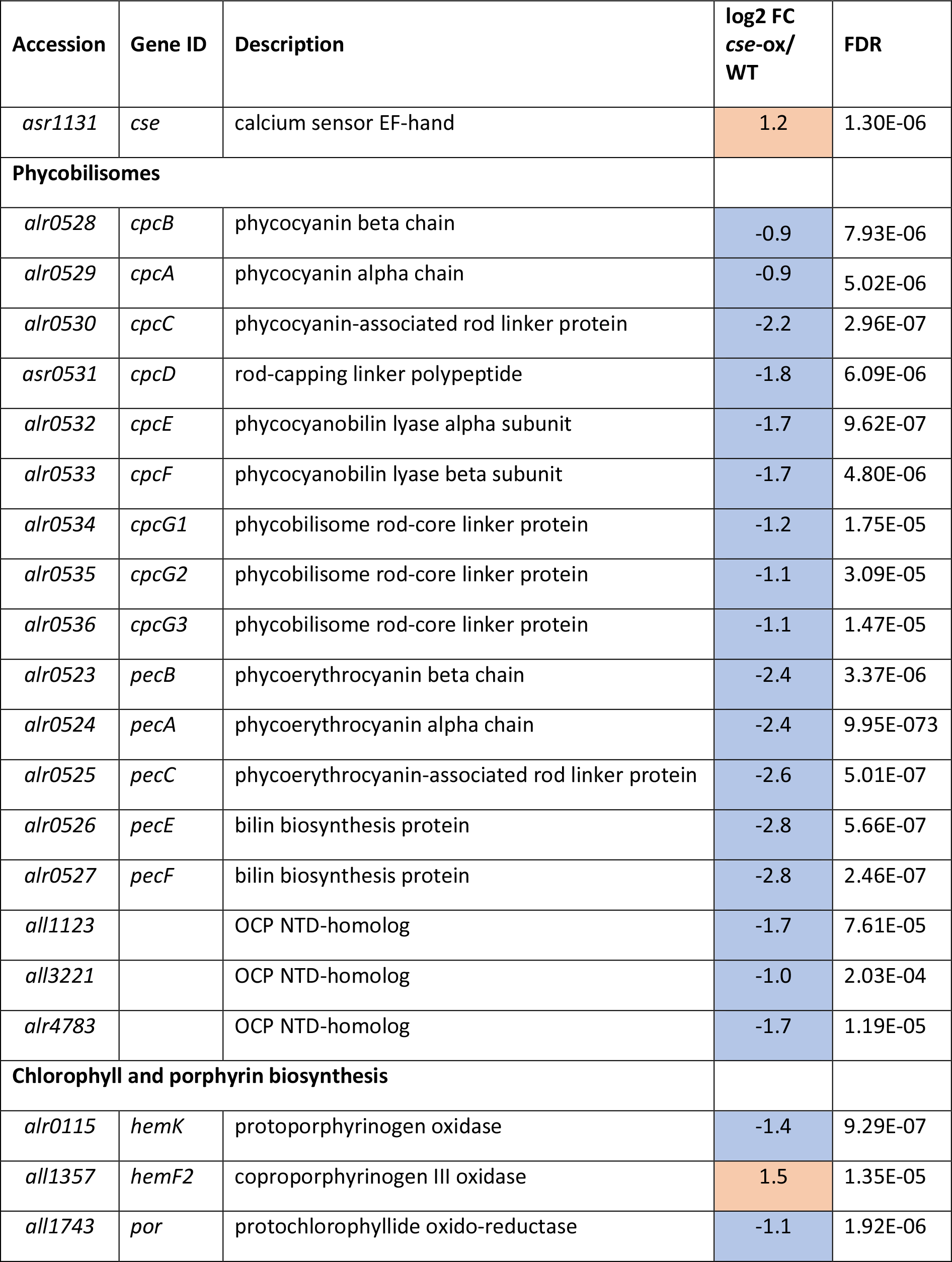
Transcription changes in the *cse*-ox mutant Differential expression of genes in *cse*-ox, in comparison to WT, grown for 2 days in BG11_A_C medium in 3% CO_2_. Genes with log2 fold change (FC) values ≥0.9 (upregulated) or ≤-0.9 (downregulated) are shown shaded red and blue, respectively. False discovery rates (FDR) were calculated using the Benjamini-Hochberg method. For the gene cluster marked with an asterisk, the average log2FC and largest FDR value are shown

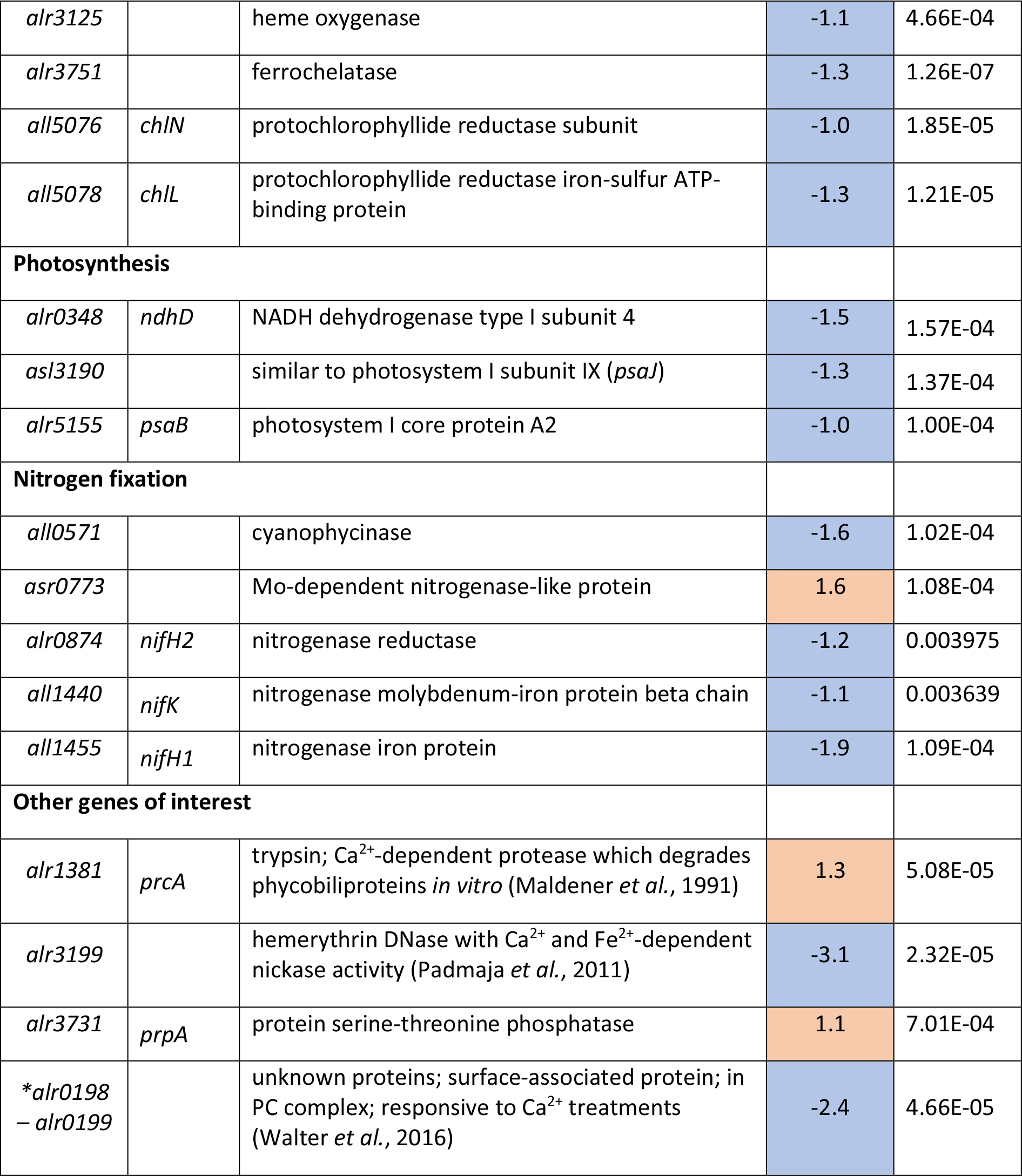

### 3.5 Photosynthetic pigment-protein complexes and photosynthetic activity are altered in cse-ox

Energetic connectivity within the photosynthetic pigment-protein complexes was studied by fluorescence emission at low temperature (77K) and room temperature in whole cell samples excited with monochromatic light of 440 nm (specific for chlorophylls) or 580 nm (specific for PBS). Chlorophyll excitation revealed a PSII:PSI ratio of about 1:4 in both the WT and *cse*-ox (Figure 4A). PBS excitation at 77K showed the peak at 646 nm, which corresponds to PC [23], to be severely decreased in *cse*-ox compared to WT, while the APC peak at 663 nm was significantly higher in the *cse*-ox (Figure 4B). At room temperature, the APC peak (660 nm) was also significantly higher in the over-expressor than in WT, while the shoulder for PC (640 nm) was significantly lower in *cse*-ox.

**Figure 4.**
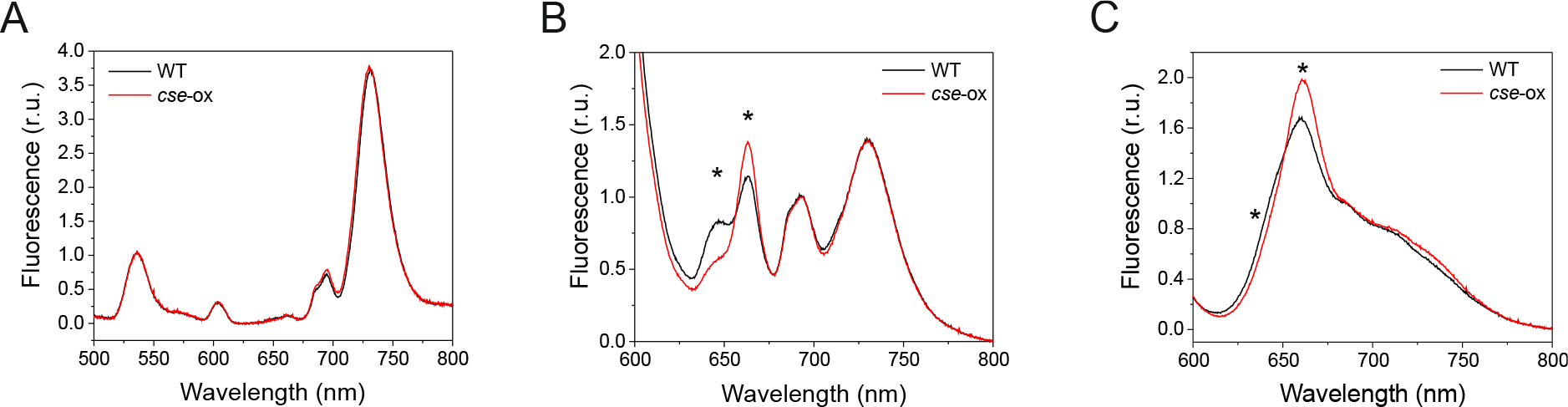
Fluorescence emission spectra of 2 day-old *Anabaena* wild-type (WT) and *cse* overexpression (*cse*-ox) strains grown in BG11_A_C **(A)** Emission at low temperature (77K) after chlorophyll excitation with 440 nm monochromatic light. Spectra are normalised to internal standard Eosin Y (5 μm, peak at 536 nm), emission peaks from PSII and PSI are located at 694 nm and 730 nm, respectively; **(B)** Emission at low temperature (77K) after phycobilisome excitation with 580 nm monochromatic light. Spectra are normalised to the PSII peak at 694 nm. Other peaks show emission from phycocyanin (646 nm), allophycocyanin (663 nm), phycobilisome terminal emitter (687 nm) and PSI (730 nm); **(C)** Emission at room temperature (25°C) after excitation of phycobilisomes with 580 nm monochromatic light. Spectra are normalised to the peak at PSII peak at 685 nm. Other peaks show emission from allophycocyanin (660 nm), phycocyanin (640 nm) and PSI (715 nm). All curves represent mean values from at least three biological replicates. Significant differences between WT and mutant samples are indicated with asterisks (*t*-test *P*<0.05)

Photosynthetic performance was monitored with a Dual-PAM-100 (Walz) on culture samples that were dark-adapted and then illuminated with constant actinic light (50 μmol photons m^−2^ s^−1^). The *cse*-ox cultures showed higher fluorescence in the absence of actinic light (F_0_ = 1.4) in comparison to the WT (F_0_ = 1.1), as well as higher steady state fluorescence under actinic light (F_s_), but saturating light pulses induced smaller F_m_’ values relative F_s_ (Figure 5A). The fluorescence signal monitored during light-to-dark transitions showed the transient rise in fluorescence that was prominent in the WT after removal of actinic light to be significantly diminished in the *cse*-ox strain (Figure 5A inset). F_v_/F_m_ determination in the presence of DCMU showed that PSII photochemical efficiency was around 65% lower in *cse*-ox in comparison to the WT value (Figure 5B), and the Y(II) parameter calculated during step-wise increases in actinic irradiance also showed that the maximum PSII quantum yield in *cse*-ox was significantly lower than in WT in light intensities from 0 to 220 μmol photons m^−2^ s^−1^, although no significant difference between the strains was observed at higher light intensities (Figure 5C). Analysis of PSII fluorescence decay after a single flash revealed significantly faster decay in *cse*-ox during the initial phase (1.5 – 3.2 ms), and slower decay in the second phase (0.006 – 3 s) (Figure 5D).

**Figure 5.**
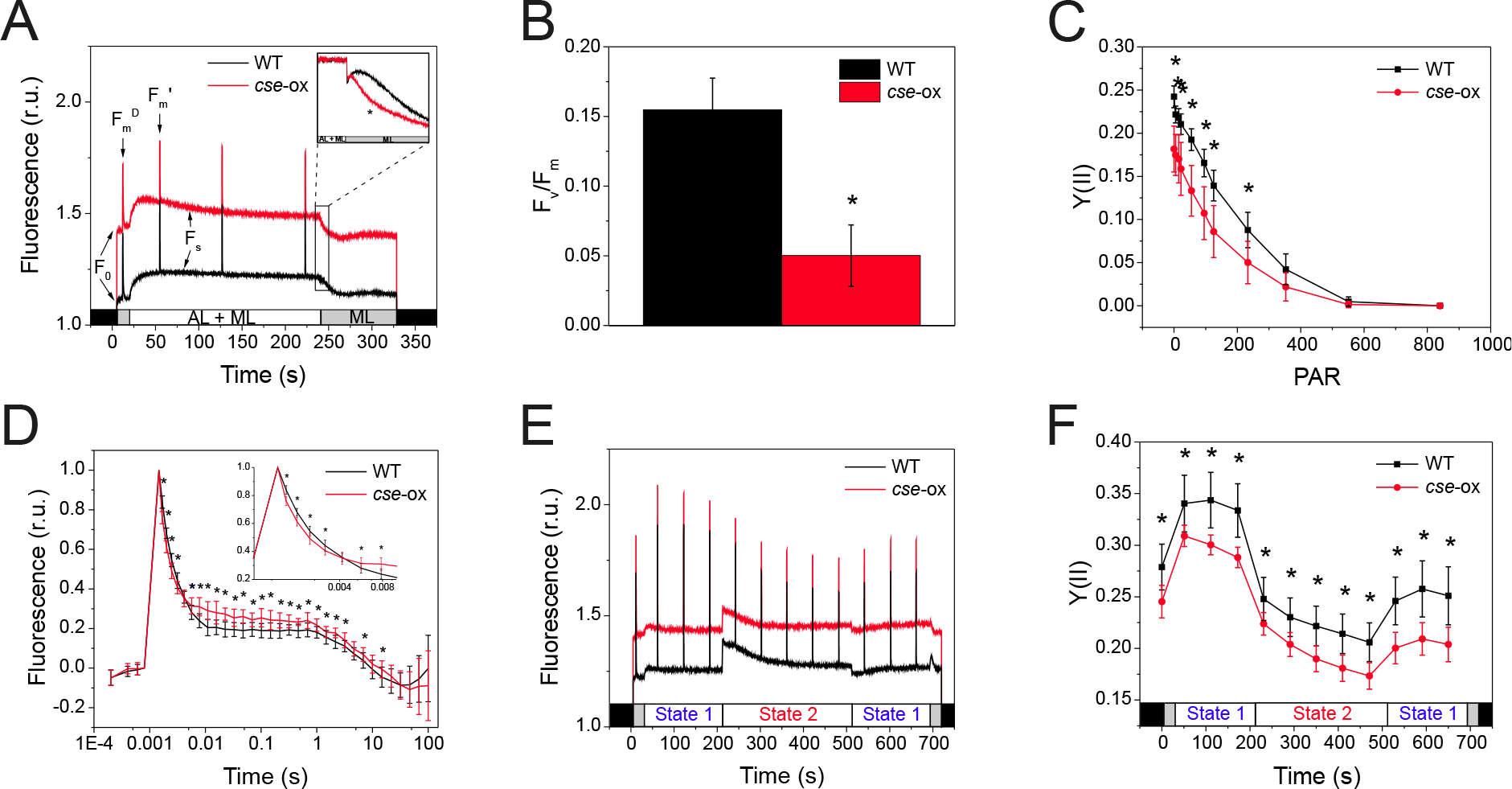
PSII-fluorescence parameters of 2 day-old *Anabaena* wild-type (WT) and *cse* overexpression (*cse*-ox) strains grown in BG11_A_C **(A)** Fluorescence induction during illumination with only measuring light (ML; 5 μmol photons m^−2^ s^−1^) or with ML + actinic light (AL; 57 μmol photons m^-2^ s^-1^). Fluorescence level under ML (F0), maximum fluorescence of open PSII reactions centres (Fm), maximum fluorescence after AL illumination (F_m’_) and steady state fluorescence under AL (Fs) are indicated. Inset: transient rise in fluorescence after switching off AL (F0 rise normalised to Fs); **(B)** F_v_/F_m_ calculated from fluorescence measurements of dark-adapted cells in the presence of 20 μM DCMU; **(C)** PSII quantum yield (Y(II)) over increasing intensities of actinic light; **(D)** Flash-induced increase and subsequent decay in fluorescence in dark-adapted WT and *cse*-ox samples; **(E)** Chlorophyll fluorescence during state transitions in dark-adapted samples illuminated with blue light (460 nm; State 1) and red light (620 nm; State 2); **(F**) PSII quantum yield (Y(II)) calculated from fluorescence induced by saturating pulses during state transitions measurements. All data represent mean values from four biological replicates, error bars show standard deviations. Significant differences between WT and mutant samples are indicated with asterisks (*t*-test *P*<0.05)

To determine whether the capacity for state transitions was affected in *cse*-ox, chlorophyll fluorescence was measured in dark-adapted samples that were illuminated with actinic blue light to induce state 1, then with actinic red light to induced state 2, and then again with blue light. The results showed equivalent capacity for state transitions in both *cse*-ox and WT, despite a higher basal fluorescence level in the over-expressor, as detected earlier (Figure 5E). Y(II) in *cse*-ox during state transition measurements was consistently and significantly lower than in the WT in both state 1 and state 2 conditions (Figure 5F).

P700 maximum quantum yield Y(I), and donor and acceptor side limitation (Y(ND) and Y(NA), respectively) were measured by pre-illuminating culture samples with far-red light, followed by illumination with red actinic light and periodic application of saturating pulses. After far red light illumination, *cse*-ox had about 20% higher Y(I) (Figure 6A) and 30% lower Y(ND) (Figure 6B), compared to WT. P_m_ measurements showed equivalent levels of P700 oxidation in both WT and over-expressor (data not shown). During excitation with red light, *cse*-ox showed significantly lower Y(I) levels and higher Y(NA) in comparison to the WT (Figures 6A and 6C). Y(I) values measured over increasing light intensities showed lower Y(I) in the *cse*-ox between 0 to 220 μmol photons m^−2^ s^−1^, with values at high light intensities equivalent to WT (Figure 6D). Absorbance of P700^+^ measured in dark-adapted cells during 5 s strong far red light illumination showed rapid increases P700 oxidation under far red in both genotypes, but also revealed a decrease in P700^+^ signal in *cse-*ox during far red illumination that was not seen in WT (Figure 6E).

**Figure 6.**
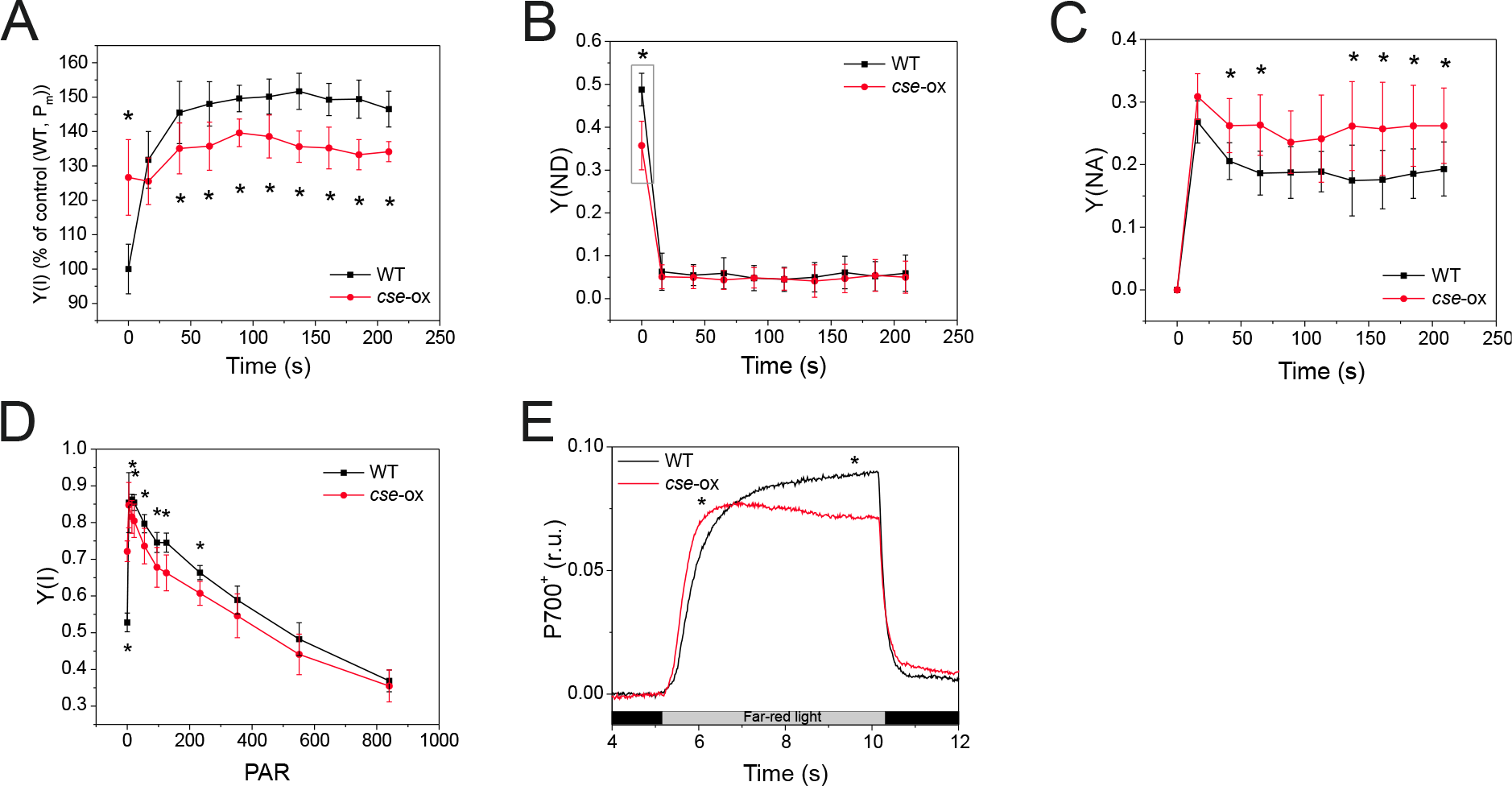
PSI parameters measured in 2 day-old *Anabaena* wild-type (WT) and *cse* over-expression (*cse*-ox) strains grown in BG11_A_C **(A)** P700 maximum quantum yield (Y(I)) in WT and *cse*-ox culture samples during illumination with red actinic light after 10 s illumination with far red light (for determination of maximum P700^+^). Y(I) values are expressed relative to WT time-point 0, which represents 100% Y(I); **(B)** Donor side limitation of PSI (Y(ND)) and **(C)** acceptor side limitation of PSI (Y(NA)) calculated from P700^+^ values measured during actinic light illumination in **(A)**; **(D)** P700 maximum quantum yield from (Y(I)) determined over increasing intensities of red actinic light; **(E)** P700 oxidation and reduction during far red illumination, normalised to baseline values only. All data represent mean values from four biological replicates, error bars show standard deviations. Significant differences between WT and mutant samples are indicated with asterisks (*t*-test *P*<0.05)

Oxygen uptake/evolution measured with a Clark-type oxygen electrode revealed about 35% lower respiration rate in *cse*-ox during 5 min dark incubation, compared to the WT control (Figure 7A), while about 20% higher net oxygen evolution rate was recorded in the over-expressor under illumination with 400 μmol photons m^−2^ s^−1^ (Figure 7B).

**Figure 7.**
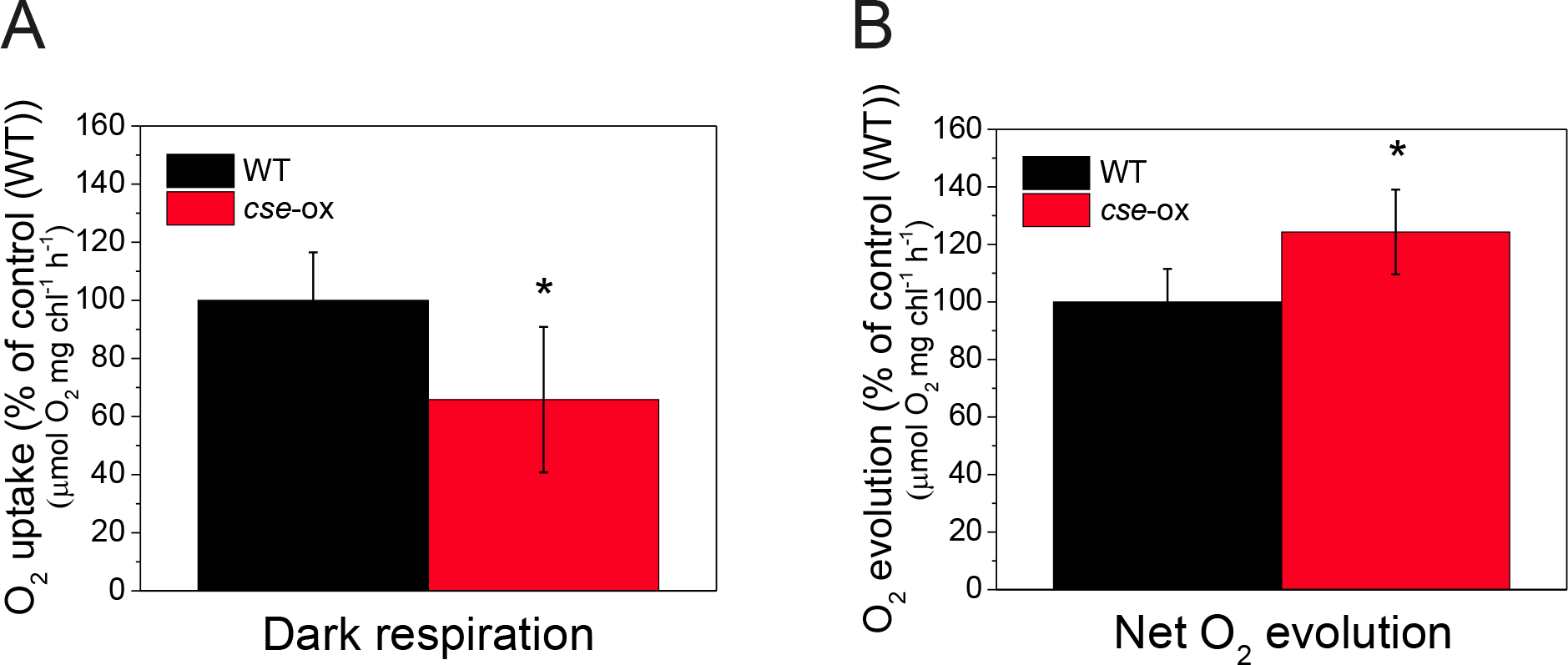
Oxygen exchange rates in 2 day-old *Anabaena* wild-type (WT) and *cse* over-expression (*cse*-ox) strains grown in BG11_A_C **(A)** Oxygen uptake in WT and *cse*-ox in darkness; **(B)** Net oxygen evolution in high light (400 μmol photons∙m^−2^·s^−1^). Values represent means of four biological replicates, error bars show standard deviations. Significant differences between WT and mutant samples are indicated with asterisks (*t*-test *P*<0.05)

### 3.6 Abundance of protein complexes in the thylakoid membrane is affected by increased cse levels

To evaluate the composition of photosynthetic protein complexes, purified thylakoid membranes were solubilised with 1.5% DM, and protein complexes were separated by large pore blue native (lpBN)-PAGE using a 3.5 - 12.5% acrylamide gradient. Blue native PAGE separation revealed substantial increases in abundance of monomeric PSI and PSII complexes in the *cse-*ox strain compared to the WT, as well as an increase in monomeric PSII lacking the CP43 protein (Figure 8A). PSII dimers were virtually absent from cse*-ox*, and PSI dimers were also less abundant, while the PSI tetramer band appeared to be more prominent in *cse*-ox compared to WT. An unknown blue band migrating in the upper region of the lpBN-PAGE gel was substantially less abundant in *cse*-ox (see black arrow in Figure 8A). In the lower region of the BN gel, several blue bands appeared to have different migrations in *cse*-ox in comparison to WT (Figure 8A).

**Figure 8.**
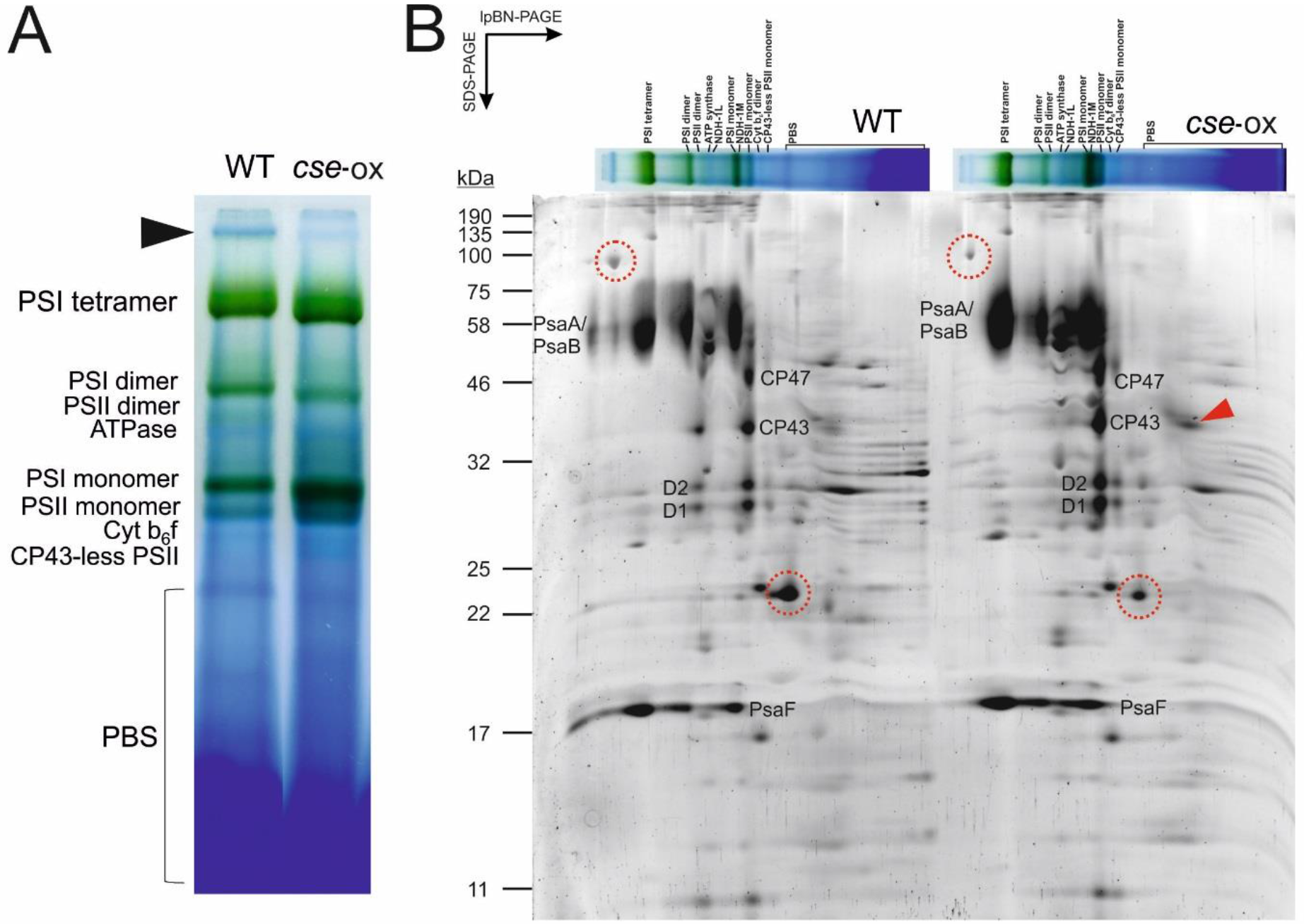
Photosynthetic thylakoid membrane protein complexes of 2 day-old *Anabaena* wild-type (WT) and *cse* over-expression (*cse*-ox) strains grown in BG11_A_C **(A)** Thylakoid membranes equivalent to 75 μg protein were solubilised with 1.5% *n*-dodecyl-β-D-maltoside (DM) and separated by large pore blue native-PAGE using an acrylamide gradient of 3.5 - 12.5%. Major protein complexes are labelled. Black arrow indicates unknown high molecular weight complex putatively associated with PSI (see text); **(B)** Separation of protein complexes in second dimension using SDS-PAGE containing 12% acrylamide. Proteins were stained with Sypro Ruby Protein Gel Stain. Characteristic subunits of PSI and PSII complexes are labelled. Red arrow indicates free CP43 protein. Protein spots cut from the WT gel are circled in red in both WT and *cse-*ox gels. Sizes of molecular weight markers are indicated on the left side

Subunit proteins of photosynthetic complexes separated by lpBN-PAGE were further separated according to their molecular mass using SDS-PAGE in the second dimension (Figure 8B). Protein spots corresponding to CP47, CP43, D1 and D2 subunits of PSII were clearly derived from both PSII dimers and monomers in WT samples, while these spots were virtually absent from positions corresponding to the larger PSII dimer complex in *cse*-ox and were more abundant in positions corresponding to the PSII monomer. PSII monomers lacking CP43 were also more abundant in *cse*-ox, as was a spot corresponding to free CP43 protein (see red arrow in Figure 8B). PsaA/B and PsaF subunits derived in denaturing PAGE from the upper region of the lpBN-PAGE gel (above the PSI tetramer band) revealed the existence of very high molecular weight PSI complexes in WT thylakoids that appeared to be absent from the *cse*-ox thylakoids. To investigate whether these PSI complexes were associated with light-harvesting antennae, a 100 kDa protein spot in the SDS-PAGE gel (see Figure 8B), which corresponded to the high molecular weight band in the native gel described above, was cut from the WT gel and investigated by mass spectrometry (MS). A second protein spot of approximately 20 kDa, which was less abundant in *cse*-ox thylakoids, and appeared to migrate with a PBS band in the lpBN-PAGE, was also identified by MS (Figure 8B). “Unknown proteins” all3041 and alr2489 were identified with high confidence in the first and second protein spots, respectively. Both proteins were previously identified in *Anabaena* thylakoids [49], although no connection between these proteins and photosynthesis was apparent.

## 4. Discussion

Internal Ca^2+^ concentration ([Ca^2+^]_i_) in cyanobacteria is tightly controlled by ion channels/transporters and Ca^2+^-binding proteins. Changes in free [Ca^2+^]_i_ regulate many cyanobacterial processes, including phototactic orientation, heterocyst differentiation and frequency, PBS degradation, hormogonia differentiation and gliding [reviewed in 50]. However, the mechanisms of Ca^2+^-sensing and signalling in cyanobacteria are poorly understood. We describe here the discovery of CSE, a previously unreported EF-hand protein that binds Ca^2+^ *in vitro* and is highly conserved in filamentous, mostly heterocystous cyanobacteria. All CSE homologs identified in cyanobacteria genome databases contain an N-terminal EF-hand that perfectly matches the EF-hand consensus sequence [46]. The C-terminal EF-hand appears to have lower affinity for Ca^2+^ due to substitutions of key residues in this domain, which was supported by thermodynamic data that indicated a single Ca^2+^-binding site (Table 1). Alterations in CSE structure upon Ca^2+^-binding detected by CD analysis (Figure 4E) resembles the Ca^2+^-mediated conformational changes of other EF-hand proteins such as calmodulin and troponin C [51], suggesting that Ca^2+^-binding may regulate an interaction between CSE and receptor proteins as part of a Ca^2+^-sensitive signalling pathway.

To investigate the function of CSE, an overexpression strain (*cse*-ox) was generated with a two-fold increase in *cse* expression compared to WT *Anabaena* (Table 3). This moderate increase in *cse* expression was sufficient to impair overall growth of the mutant strain, as well as causing yellow colouration of cultures and decreased levels of phycocyanobilin (PCB) pigment (Figure 3), which can indicate N deficiency and subsequent degradation of N-containing pigments [53]. A role for CSE in N metabolism was also alluded to by almost exclusive occurrence of *cse* gene homologues in filamentous, N-fixing cyanobacteria. Considering these findings together with previously established links between Ca^2+^ signalling and N step-down [30,53,54], we investigated whether the growth penalty in *cse-*ox may be due to impaired N metabolism. However, heterocyst frequency and nitrogenase activity in cells grown in media lacking combined N sources did not differ significantly from WT levels (Table 2). Instead, the observed decrease in growth was consistent with a decrease in photosynthetic activity and substantial alterations in the composition of photosynthetic pigment-protein complexes, including the phycobilisome (PBS) light-harvesting antennae, and the multimeric states of PSII and PSI complexes. Notably, we also created knockout mutant *Anabaena* strains lacking the *cse* gene, which displayed many of the same phenotypical characteristics displayed by the *cse*-ox strain, including slow growth, altered photosynthetic activity and thylakoid membrane organisation. However, substantial variability was observed between different clones of the ∆*cse* knockout, in contrast to the phenotype of *cse*-ox that was consistent between several individual clones. Therefore, only the *cse*-ox strain is described here. Nonetheless, striking similarity between cells with moderate over-abundance of CSE and those lacking CSE supports the role of the CSE protein, and proper regulation of Ca^2+^, as the underlying factor of the phenotypes described here, rather than possible artefactual causes related to the genetic modifications.

Transcriptomic analysis of *cse*-ox revealed strong downregulation of gene clusters encoding protein components of the light-harvesting complexes, in comparison to the WT. The entire *pec* gene cluster, which encodes the phycoerytherocyanin (PEC) proteins, and the *cpc* gene cluster, which encodes the phycocyanin (PC) proteins, were the most strongly downregulated genes in the mutant (Table 3). An exception was *cpcG4*, which is transcribed independently from the *cpc* cluster [20,27]. PC and PEC proteins make up the phycobiliprotein rods that radiate from the allophycocyanin (APC) core cylinders of the PBS complex. Unlike the peripheral PBS subunits, expression of genes encoding APC subunits was not affected in *cse*-ox, suggesting that PBS rods in *cse*-ox may be truncated. Disturbed PBS function in *cse*-ox was also apparent in fluorescence emission measurements at 77K and room temperature, both of which showed substantially lower emission from PC at 646 nm, and increased emission from APC at 663 nm, after PBS excitation with 580 nm light (Figure 4). Light energy absorbed by the PBS is normally transferred sequentially from PEC to PC, to APC, then to the terminal emitter ApcE and finally to P680 (PSII reaction centre chlorophyll) to initiate linear electron transport (LET) towards PSI [22]. Reduced PC fluorescence emission was consistent with lower PC and PEC abundance, while enhanced fluorescence emission from APC (Figure 4) indicates inefficient energy transfer between PBS and the photosystem reaction centres, resulting in release of excitation directly from APC. Increased APC fluorescence has previously been linked to abnormal PBS rod composition [24].

PSII:PSI ratios were equivalent between WT and the *cse*-ox mutant (Figure 4A), indicating that defective excitation transfer from PBS was not due to any changes in photosystem stoichiometry in the mutant. On the other hand, the virtual absence of PSII complex dimers (Figure 8A), which are required for PSII-PBS connectivity [24,55–59], most likely had a substantial negative impact on normal excitation transfer from PBS. Corresponding increases in PSII monomers, both with and without the CP43 subunit, as well as free CP43 protein, were clearly evident in *cse*-ox thylakoids (Figure 8). This finding indicates a high rate of PSII damage and turn-over in the mutant [60,61], which may be caused by over-reduction of the photosynthetic electron carriers and increased excitation pressure on PSII. Indeed, such over-reduction was evident in high levels of chlorophyll *a* fluorescence indicative of more closed PSII centres in both darkness and actinic light, and in the slower rate of fluorescence decay phase that is attributed to slower binding of oxidised PQ to the Q_B_ pocket (see Figure 5D; 0.006 – 3 s). A steady decrease in PSII efficiency observed in *cse*-ox exposed to “state 1 light” (Figure 5F) may have been caused by progressive PSII damage under light conditions that induce PBS association with PSII. Oddly, PSII damage may have been exacerbated in the mutant by downregulated expression of genes encoding three helical carotenoid proteins (HCPs) that are homologous to the OCP N-terminal domain [62]. Two of the downregulated HCPs are active in quenching of singlet oxygen (^1^O_2_) [63], suggesting a decreased ^1^O_2_ scavenging capacity that may have enhanced the level of PSII damage in *cse*-ox. The conventional OCP gene (*all3149*) that quenches ^1^O_2_ as well as excess excitation directly from PBS under light stress [64,65], was not found to be differentially expressed in our data.

Altered PSI activity was detected in *cse*-ox, including re-reduction of oxidised P700^+^ during far-red (FR) light illumination, lower Y(ND) after FR exposure, and higher Y(NA) under actinic light (Figure 5). These effects can also be attributed to over-reduction of photosynthetic electron carriers in the mutant leading to an abnormally high flow of electrons to the PSI donor side. Clearly this over-reduction could not be alleviated by PSI activity, in spite of an apparent increase in PSI complex abundance evident in *cse*-ox thylakoids (Figure 8). It is possible that inefficient excitation transfer to PSI from abnormal PBS lacking PC and PEC (discussed above) may have inhibited PSI electron transport (Figure 8).

Inhibited respiration in *cse*-ox was evident in the lower rate of oxygen uptake in darkness (Figure 7A), and in the higher rate of net oxygen evolution in the light (Figure 7B). Considering the lower PSII activity in *cse*-ox (described above), higher net oxygen evolution is unlikely to have been caused by increased PSII water-splitting, instead implying lower oxygen consumption in the over-expressor in the light. Respiration in cyanobacteria provides several alternative pathways of electron transport from metabolites, via PQ, to terminal oxidases in the thylakoid and plasma membranes, operating in both light and darkness [reviewed in 66]. Decreased electron flux through respiratory pathways could explain over-reduction of photosynthetic electron carriers and the subsequent negative effects on PSII and PSI activity described above. Indeed, higher chlorophyll fluorescence and over-reduced P700 were observed in mutant cyanobacteria lacking the respiratory terminal oxidase (RTO) enzymes cytochrome *bd* quinol oxidase (Cyd) and cytochrome *c* oxidase (COX) [67]. The results observed in *cse-*ox could also be attributed to decreased oxygen photoreduction by the flavodiiron proteins, which are especially important for protection of PSII and PSI in stress conditions [68].

## 5. Conclusions and perspectives

The results presented here show the newly discovered CSE protein to be a *bona fide* Ca^2+^-binding EF-hand protein. Therefore, disrupted electron transport and thylakoid protein complex formation/stability in *Anabaena* with over-abundance of CSE is thought to relate to modified Ca^2+^ signalling and/or a change in Ca^2+^ homeostasis. Although the precise mechanism behind the observed phenotypes of *cse*-ox has not yet been identified, several factors that regulate photosynthetic energy balance in cyanobacteria have established links with Ca^2+^. The flavodiiron protein Flv4 contains Ca^2+^-binding sites, and the presence of cations (Ca^2+^ or Mg^2+^) was shown to influence partitioning of the Flv2-Flv4 dimer between the thylakoid membrane (cations present) and the soluble fraction (cations absent) [69]. A mutant strain lacking *flv4* showed substantial phenotypic overlap with the *cse*-ox mutant, including increased PSII monomer:dimer ratio, over-reduction of the PQ pool and higher fluorescence emission from detached APC [69–71]. Furthermore, expression of *flv4* in *Anabaena* was downregulated by a shift to low Ca^2+^ conditions [30]. It is possible that over-expression of *cse* may interfere with Flv4 association with the thylakoid, leading to PQ over-reduction and increased PSII monomerisation. A Ca^2+^-binding site is conserved in prokaryotic and eukaryotic orthologues of subunit A of the aa3-type COX terminal oxidase [72–74]. This site appears to be also present in the predicted *Anabaena* COXA structure (see supplemental Figure S4). COX activity was downregulated by Ca^2+^-binding to mitochondrial COXA [75], suggesting that abnormally high or otherwise unregulated Ca^2+^ levels in *cse*-ox may have led to decreased respiration activity and subsequent over-reduction of PQ through inhibition of COX [67].

The Ca^2+^-induced conformational change in CSE, taken together with the activity of other EF-hand proteins such as calmodulin and troponin C, suggests that CSE is likely to interact with partner proteins in a Ca^2+^-dependent manner. Interaction partners may include protein phosphatases, such as PrpA which was upregulated in *cse*-ox, or protein kinases. PBS proteins are known to be phosphorylated when assembled [76,77], and so dysfunctional Ca^2+^-dependent regulation of PBS phosphorylation may explain the modified PBS composition identified here. Furthermore, abnormally-abundant proteins in *cse*-ox identified by MS (all3041 and alr2489) were found to be heavily phosphorylated in the WT (data not shown). Identification of interaction partners will be an important step in understanding the molecular mechanism of the CSE protein.

An outstanding question relates to the evolution of CSE, which appears almost exclusively in filamentous, N-fixing cyanobacteria. Changes in [Ca^2+^]_i_ in *Anabaena* have been linked to the differentiation of vegetative cells into heterocysts under N-deficient conditions [48,53], and appear to play a role in communicating C:N homeostasis [30]. The current work shows that CSE over-abundance does not affect normal heterocyst differentiation and nitrogenase activity; however, the observed effects on photosynthetic activity may be expected to downregulate carbon metabolism and thereby impact C:N balance. During heterocyst differentiation, substantial changes to the organisation and function of thylakoid membranes are known to occur, including decreased abundance of PBS [78] and increased respiration via COX [79,80], which may link the photosynthetic phenotypes reported here with disrupted Ca^2+^ signalling in filamentous strains.

## 6. Acknowledgements

J.W. and P.J.G thank Ilaria Mancini for running lpBN-PAGE gels, Andrea Trotta and Azfar Ali Bajwa for protein MS analysis, Yagut Allahverdiyeva, Natalia Battchikova and Mikko Tikkanen for help with data analysis and useful discussions. K.A.S. thanks Marcus Hartmann and Reinhard Albrecht for the assistance with CD spectroscopy. Financial support was provided from EU Marie Curie ITN CALIPSO project GA ITN 2013–607607 (J.W.) and Academy of Finland projects 307335 and 303757 (E-M.A.) and 26080341 (P.J.G.). K.A.S and K.F. were supported by Deutsche Forschungsgemeinschaft (DFG).

## Supplementary figures and tables

**Figure S1:**
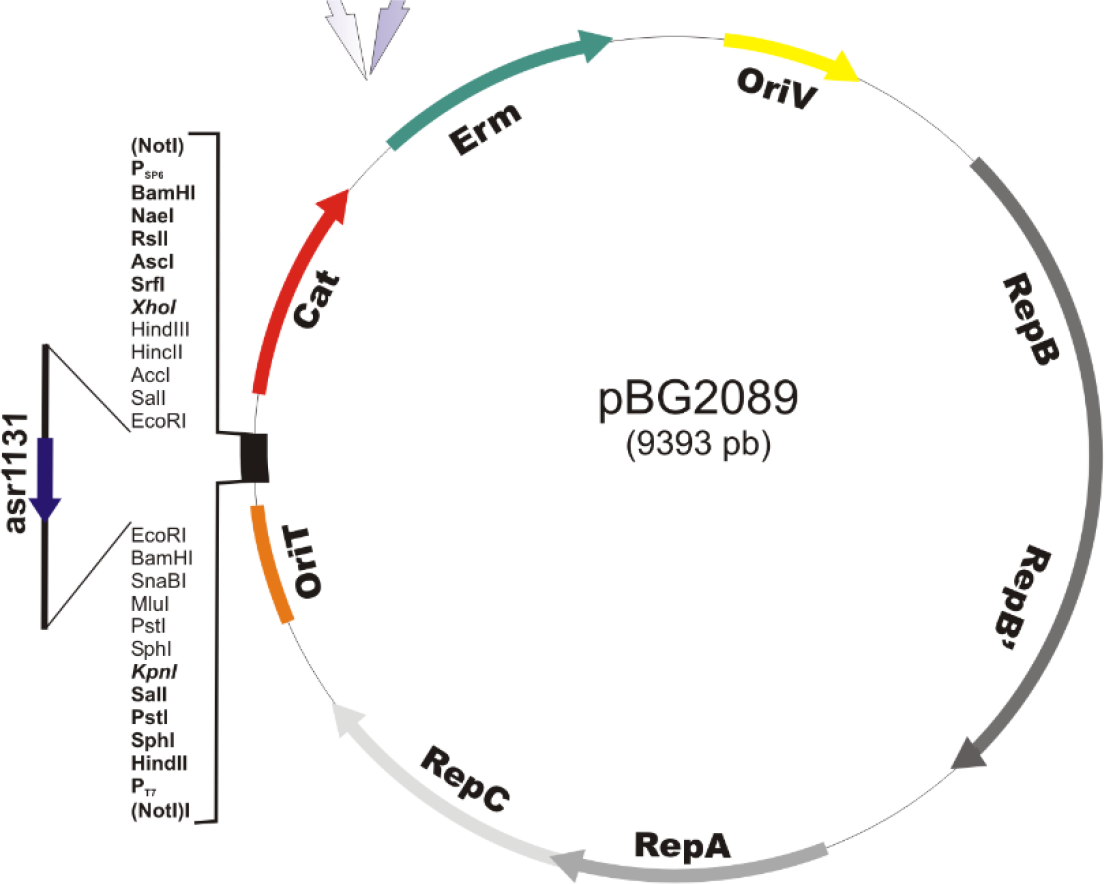
*cse* overexpression vector *pBG2089* The *asr1131* gene under the control of its native promoter was cloned into the plasmid *pRL1342*, a *RSF1010* derivative designed for very low copy numbers, generating plasmid *pBG2089*.

**Figure S2:**
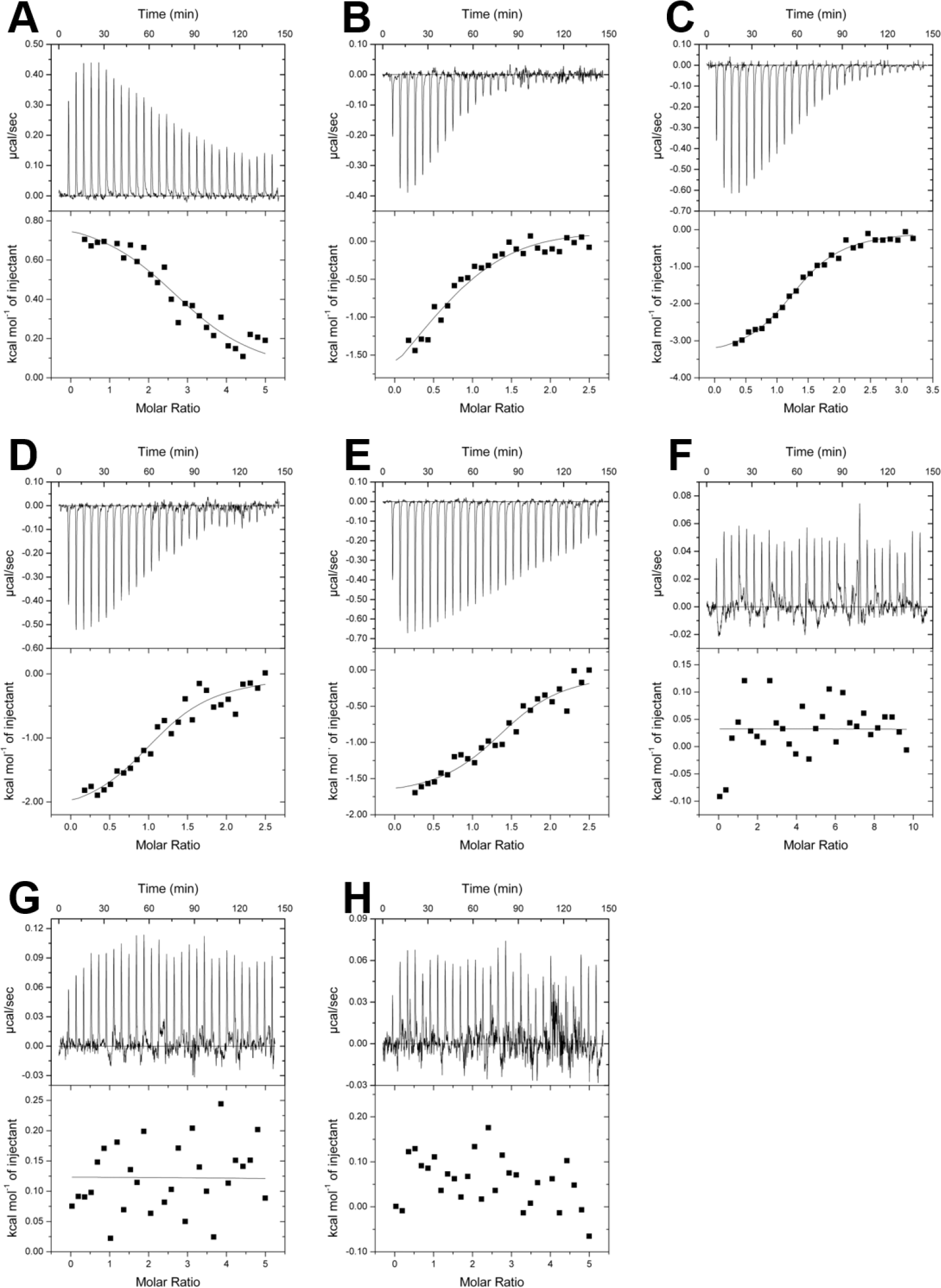
Isothermal titration calorimetry (ITC) metal-binding assay of recombinant His-tagged CSE in different buffers. **A.** 85 μM CSE in 20 mM Tris buffer against 2 mM MgCl_2_ **B.** 85 μM CSE in 20 mM Tris buffer + 100 mM NaCl against 1 mM CaCl_2_ **C.** 66 μM CSE in 20 mM Tris buffer + 300 μM MgCl_2_ against 0.75 mM CaCl_2_ **D.** 85 μM CSE in 20 mM Tris buffer + 300 μM MgCl_2_ + 100 mM NaCl against 1 mM CaCl_2_ **E.** 85 μM CSE in 20 mM Tris buffer + 2 mM MgCl_2_ + 100 mM NaCl against 1 mM CaCl_2_ **F.**66 μM CSE in 20 mM Tris buffer + 150 μM CaCl_2_ against 3 mM MgCl_2_ **G.** 85 μM CSE in 20 mM Tris buffer + 100 mM NaCl against 2 mM MgCl_2_ **H.** 85 μM CSE in 20 mM Tris buffer + 100 mM NaCl + 150 μM CaCl_2_ against 2 mM MgCl_2_

**Figure S3:**
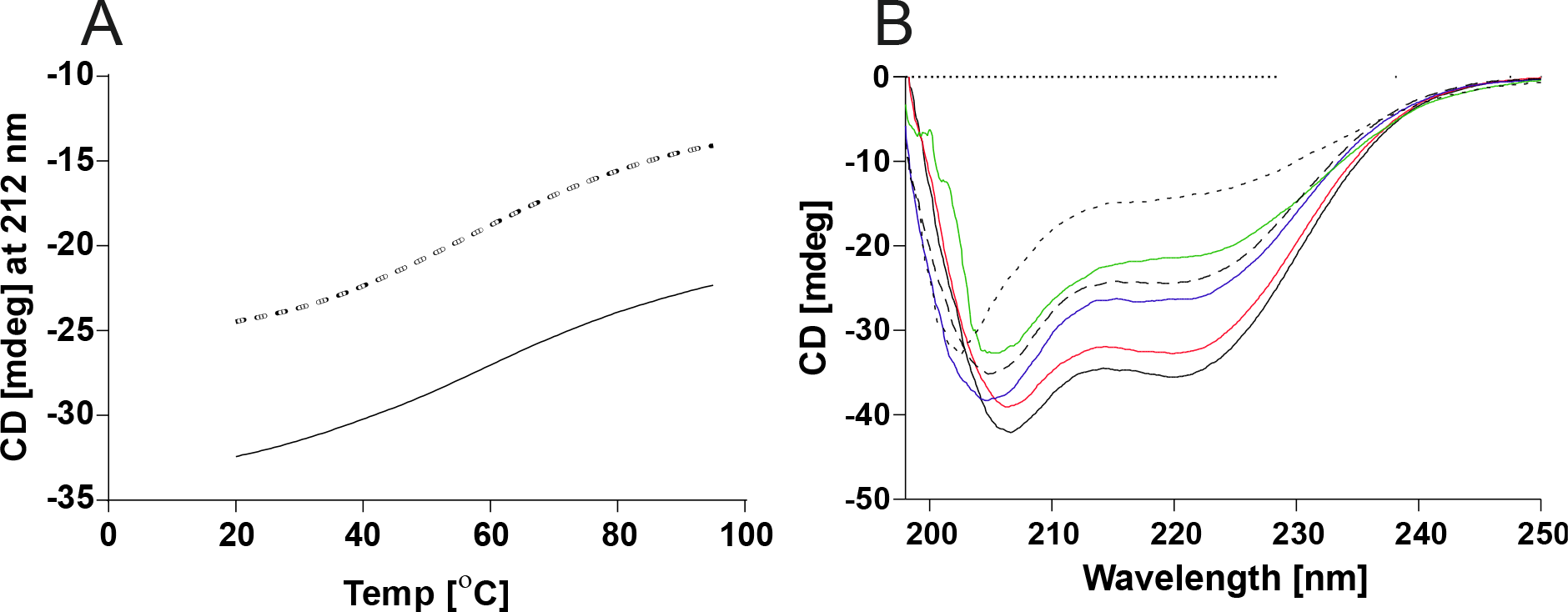
Circular dichroism (CD) spectra of recombinant His-tagged CSE in different buffers. **A.** CSE melting curves from 20-95°C at 212 nm in 20 mM Tris buffer (dashed line) and 20 mM Tris buffer + 100 mM NaCl + 1 mM CaCl_2_ (solid line) **B.** CD spectra of CSE in 20 mM Tris buffer at 20°C (after melting; blue solid line), at 20°C (black dashed line), at 95°C (black dotted line), and in 20 mM Tris buffer + 100 mM NaCl + 1 mM CaCl_2_ at 20°C (after melting; black solid line), at 20°C (red solid line), at 95°C (green solid line)

**Figure S4:**
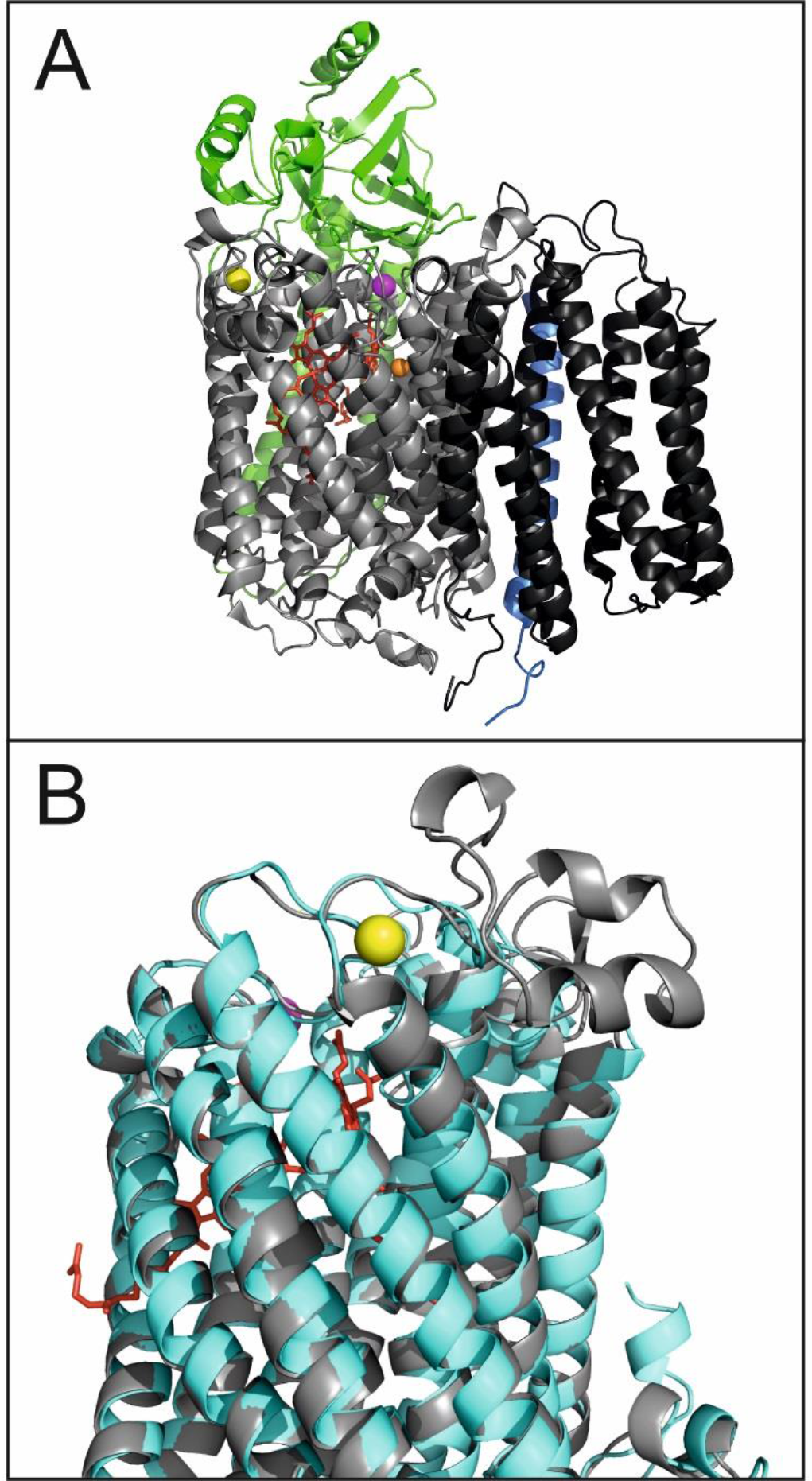
Calcium-binding in cytochrome *c* oxidase subunit COXA **A.** Structure of cytochrome c oxidase from Rhodobactor sphaeroides (PBD 1M56) [74], showing subunits COXA (light grey), COXB (green), COXC (dark grey), COXD (blue). Cofactors shown are Ca^2+^ (yellow sphere), Cu^2+^ (orange sphere), Mg^2+^ (magenta sphere) and haem (red sticks); **B.** Predicted protein structure of *Anabaena* sp. PCC 7120 COXA (alr0951; cyan) aligned with *R. sphaeroides* COXA subunit (grey; same as in **A**), to highlight putative Ca^2+^-binding site in *Anabaena* COXA.

**Table S1.**
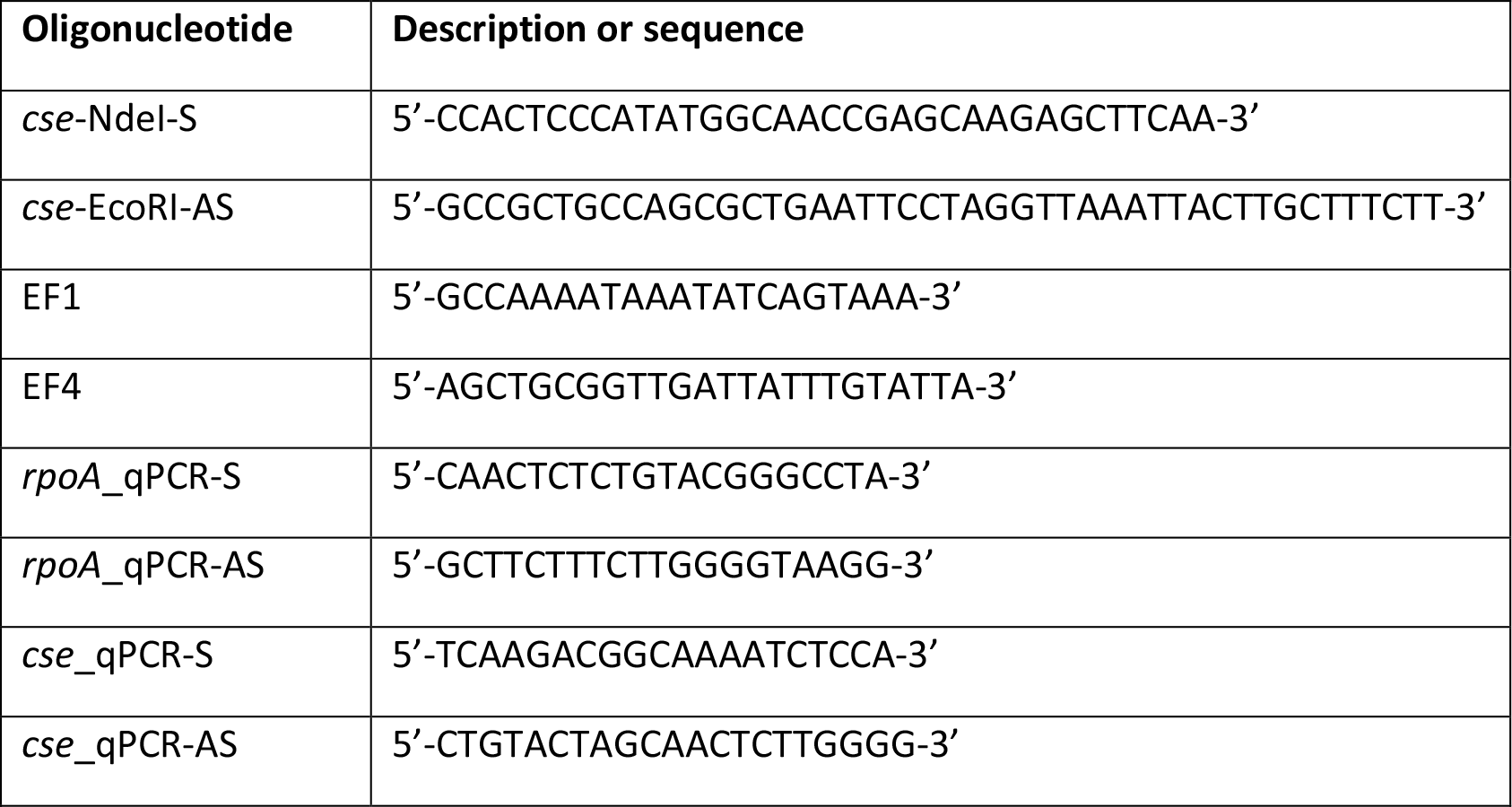
List of oligonucleotides used in the current study

